# Multilevel regulation of the *glass* locus during *Drosophila* eye development

**DOI:** 10.1101/537118

**Authors:** Cornelia Fritsch, F. Javier Bernardo-Garcia, Tim-Henning Humberg, Sara Miellet, Silvia Almeida, Armin Huber, Simon G. Sprecher

## Abstract

Development of eye tissue is initiated by a conserved set of transcripton factors termed retinal determination network (RDN). In the fruit fly *Drosophila melanogaster,* the zinc-finger transcription factor Glass acts directly downstream of the RDN to control idendity of photoreceptor as well as non-photoreceptors cells. Tight control of spatial and temporal gene expression is a critical feature during development, cell-fate determination as well as maintainance of differentiated tissues. The molecular mechanisms that control expression of *glass*, however remain largely unknown. We here identify complex regulatory mechanisms controlling expression of the *glass* locus. All information to recapitulate *glass* expression are contained in a compact 5.2 kb cis-acting genomic element by combining different cell-type specific and general enhancers with repressor elements. Moreover, the immature RNA of the locus contains an alterantive small open reading frame (smORF) upstream of the actual glass translation start, resulting in a small peptide instead of the three possible glass protein isoforms. CRISPR/Cas9-based mutagenesis shows that the smORF is not required for the formation of functioning photoreceptors, but to attenuate effects of *glass* misexpression. Furthermore, editing the genome to generate *glass* loci eliminating either one or two isoforms shows that only one of the three proteins is critical for formation of functioning photoreceptors, while removing the two other isoforms did not cause defects in developmental or photoreceptor function. Our results show that eye development and function is surprisingly robust and appears buffered to targeted manipulations of critical features of the *glass* transcript, suggesting a strong selection pressure to allow the formation of a functioning eye.

## INTRODUCTION

While genes of the retinal determination network (RDN) are necessary and sufficient for inducing eye tissue in the imaginal-disc, distinct transcription factors are subsequently involved in promoting the developmental program of cell fate determination as well as terminal differentiaton. The zinc-finger transcription factor Glass provides a critical link betwee the RDN and terminal differentiaton. Glass is required during eye development for the differentiation of photoreceptor neurons, patterning of the ommatidia, as well as for the differentiation of cone- and pigment-cells [1-4]. *glass* mutants were first discovered by H.J. Muller in 1918 and O. L. Mohr in 1919, and were named after their smaller eyes with smooth, glassy surface and altered pigmentation [5]. While it was initially assumed that photoreceptor precursors undergo apoptosis in *glass* mutants, we recently showed that these cells adapt a neuronal cell fate, extend axons and form synapses, but fail to express *rhodopsins* as well as phototransduction genes. For the determination of photoreceptor idendity, glass promotes the terminal differentiation gene *hazy* [1, 6]. Intrestingly glass acts in conjunction with distinct transcripton factors to coordinate different cell fates during eye formation. For the specification of cone cells glass acts together with d*Pax2, eyes* absent and *lozenge*, while for the formation of pigment cells it requires *escargot* [4]. Thus, dependent on the cellular context Glass is likely to control distinct developmental programs. However, mechanisms that act to control expression of glass remain largely unknown.

We here provide insight into surprisingly diverse regulatory mechanisms acting to regulate the *glass* locus. By further dissectioning a previously identified 5.2 kb genomic element we identified a set of regulatory core elements, including a general promotor, two pan-PR enhancer elements, a reciprocal enhancer element for non-PR cells, an element driving expression in a subset of PRs as well as an ocelli-specific enhancer element. By analysing a GFP reporter including the 5’UTR we identified an alterantive small open reading frame (smORF) upstream of the actual Glass translation start, resulting in a small peptide instead of the Glass protein. Interestingly editing the corresponding genomic sequences to mutate the smORF did not cause any developmental defect nor inference with physiological response of the retina nor photoattraction behaviour. However, when misexpressessing Glass in a transcript including the smORF it attenuates developmental deficits, suggesting that while evolutionarily conserved within Drosophilids the smORF is not essential for eye development, but may act to buffer Glass expression level. Moreover, we assessed the requirement and functionality of the three Glass protein isoforms by CRISPR-mediated genome editing introducing deletions into the *glass* locus resulting in the loss of one or two of the three isoforms. We analyzed these isoform mutants for the morphology of their eyes, the expression of photoreceptor markers that depend on Glass function, photoreceptor activity, and light preference behavior. We found that the short Glass PB isoform is not able to confer normal eye development and function resulting in a *glass* mutant phenotype, while the Glass PA isoform alone is fully functional. Our results suggest that the expression of *glass* is tightly regulated as the development of a functional tissue is surprisingly robust resulting in no detectable change in the physiological response or alteration in photoattraction behaviour upon deletion of the smORF. Similarly, only one of the isoforms is critical for eye development, suggesting that the other isoforms may function in a similar way to control levels of the functional protein. Since sequence comparison to closely related species show conservation of these features, such mechanisms may function to fine-tune gene expression.

## RESULTS

### An overlapping upstream open reading frame inhibits the expression of a *glass* reporter

In the developing eye, expression of *glass* is initated at the morphogenetic furrow in the eye-imaginal disc of third instar larvae and is detectable in the nuclei of all cells posterior to the morphogenetic furrow [7]. The same expression pattern is obtained with a reporter construct containing a 5.2 kb DNA fragment upstream of *glass* [8], spaning from −4190 bp to the AUG at +960 (Fig. 1A) [8]. Suprisingly, using this 5.2 kb upstream genomic sequence to drive a GFP reporter we observed GFP expression was barely detectable in the eye imaginal discs (Fig. 1B, B’’). By increasing the gain at the confocal microscope, we were able to detect a weak GFP signal posterior to the morphogenetic furrow, barely above background level (Fig. 1B’’’, B’’’’). A closer inspection of our reporter construct revealed the presence of two potential start codons in the 5’UTR of *glass*, that were also present in the GFP reporter construct, one at position +889 relative to the predicted transcription start, the other at position +955. Translation from the first start codon, if functional, may compete with the GFP start codon thus generating a protein that overlaps, but is not in frame, with the *GFP* coding sequence, resulting in the production of a 316 amino acid long protein (Fig. 1B’).

**Figure 1:**
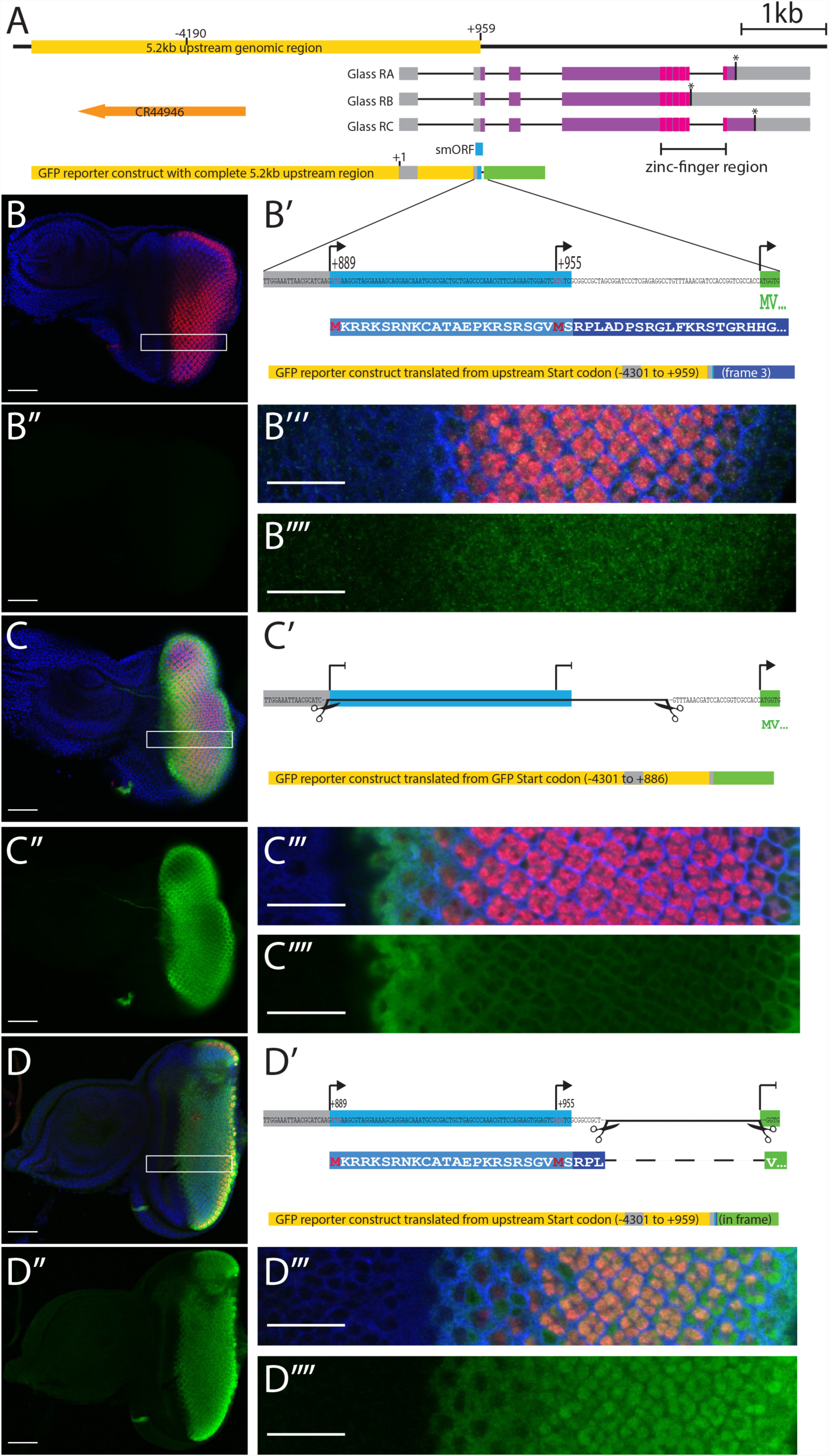
*glass* reporter constructs. A: genomic region of *glass* including the 5.2 kb regulatory region (yellow), the three glass isoforms (RA, RB, RC) with their intro-exon structures, stop codons (asterisks), the protein coding regions (purple), and the positions of the five zinc fingers (magenta). The position of the upstream overlapping open reading frame (smORF) is indicated in blue. A non-coding RNA (orange) is located upstream of the *glass* gene. The 5.2 kb upstream genomic region including the non coding exon 1 and the 5’end of exon 2 were cloned in front of eGFP (green). B: eye imaginal disc of a fly transgenic for the *glass*-GFP reporter construct (−4301 to +959). The GFP expression level is very low. B’: sequence fragment of the *glass*-GFP reporter construct including the 5’end of exon 2, the linker, and the first two codons of GFP (MV). The positions of the two upstream start codons and the GFP start codon are indicated by arrows. Translation from the upstream start codon results in the production of a protein encoded by the 3^rd^ reading frame of eGFP (first 44 amino acids shown in blue box). B’’: GFP channel alone of the disc shown in panel B. No GFP is detectable (gain: 621.7). B’’’: close up of the region indicated by the white box in B. B’’’’: GFP channel alone of the region outlined in panel B imaged with a higher gain (827.1). GFP levels are slightly higher in the posterior region of the dics. C: eye disc of a transgenic fly expressing a *glass*-GFP reporter construct in which the two upstream start codons were deleted. GFP is expressed at high level in the posterior region of the disc that will give rise to the adult eye. C’: Sequence fragment of the GFP reporter construct indicating the part that was excised to remove the upstream start codons (−4301 to +886). Translation can only start at the GFP start codon. C’’: GFP channel alone of the disc shown in panel C (gain: 621.7). C’’’: close up of the region indicated by the white box in C. C’’’’: GFP channel alone of the region outlined in panel C (gain 621.7). eGFP is mainly cytoplasmic. D: eye disc of a transgenic fly expressing a *glass*-GFP reporter construct in which the GFP start codon was deleted and the GFP coding sequence is in frame with the upstream start codon(s). D’ Sequence fragment of the GFP reporter construct indicating the part that was excised to remove the GFP start codon (−4301 to +959 in frame). The N-terminus of the resulting fusion protein between the upstream translation product and GFP is shown below). D’’: GFP channel alone of the disc shown in panel D (gain: 757.2). D’’’: close up of the region indicated by the white box in D. D’’’’: GFP channel alone of the region outlined in panle D (gain 757.2). GFP shows nuclear localization. All discs are oriented with the posterior to the right. Discs were stained with antibodies against GFP (green), Elav (red), FasIII (blue in panels B’’’, C’’’ and D’’’) and with DAPI (blue in panels B, C and D). Scale bars in panels B, B’’, C, C’’, D and D’’ are 50 µm, in panels B’’’, B’’’’, C’’’, C’’’’, D’’’ and D’’’’ are 20 µm long.

To test whether translation of GFP in our reporter construct was affected by the presence of the upstream start codon(s), we generated two additonal reporter constructs: one, in which the potential upstream start codons were deleted (Fig. 1C’), and another, in which the GFP start codon was deleted and the GFP coding sequence was brought into frame with the upstream start codons (Fig. 1D’). Both GFP reporter variants resulted in strong GFP expression posterior of the morphogenetic furrow (Fig. 1C, C’’, D, D’’). Thus, the reduced GFP expression observed in the original reporter construct was caused by the translation of the reporter construct in a different reading frame due to the presence of additional start codons upstream of the GFP coding sequence. Since the GFP reporter we used does not contain a nuclear localization signal, GFP produced from its own start codon, as in construct C’, is mainly localized in the cytoplasm (Fig. 1C’’’, C’’’’). However, when GFP was fused in frame with the smORF, it showed strong nuclear localization (Fig. 1D’’’, D’’’’), suggesting that the first 24 amino acids added to the GFP coding sequence contain a nuclear localization signal. Indeed, amino acids 2 to 20 of this fusion protein are predicted to affect nuclear localization [9]. Thus, the translation of this fusion protein starts at the first AUG codon at position +889.

### Defined enhancer elements confine cell type specificity and temporal restricted expression

In order to understand the cis-regulatory logic of *glass* expression we further dissected this genomic region in the construct that does not contain the two upstream start codons (Fig. 1C’). Using a number of restriction sites located in the upstream regulatory sequence, we generated truncations of our GFP reporter, similar to those used by Liu et al. [8], and also tested some deletions within this upstream sequence (Fig. 2A). After deleting half of the 5.2 kb fragment (construct B: −1885 to +886), GFP expression is still restricted to the region posterior of the morphogenetic furrow (Fig. 2B, B’). Further deletion of a small fragment between the BamHI and EcoRI sites (construct C: −1598 to +886) shows patchy GFP expression in the developing photoreceptor precursors (Fig. 2C, C’). While construct B is expressed in all cell types forming the presumptive eye, the expression of construct C is restricted to presumptive photoreceptor cells with variable expression levels (Fig. S1A), suggesting that the fragment from −1885 to −1598 might contain some non-photoreceptor specific enhancer. A fragment truncated at the XbaI site (construct D: −703 to +886) is expressed in all the photoreceptor precursors posterior of the morphogenetic furrow with the highest levels directly after the furrow and reduced levels towards the posterior end (Fig. 2D. This construct also shows ectopic expression in a stripe anterior of the furrow (Fig. D’ arrowhead). This misexpression of GFP is spreading over the entire eye-antenna-disc in a construct starting at the XhoI site (construct E: −239 to +886, Fig. 2E, E’), suggesting that this fragment contains a minimal promoter whose activation is independent of eye specific enhancers. We used this minimal promoter region in combination with other fragments of the enhancer to analyse the expression patterns conferred by these 5’ enhancer elements. We tested the 287 bp fragment between the BamHI and EcoRI sites that we suspected to drive expression specifically in non-photoreceptor cells based on the different expression patterns between constructs B and C. We found that this small fragment in combination with the minimal promoter (construct F: −1885 to −1598 / −239 to +886) can restrict GFP expression to the region posterior to the morphogenetic furrow (Fig. 2F). With this enhancer fragment, the GFP signal is absent in the presumptive photoreceptor cells and restricted to the cells surrounding the photoreceptor precursors (Fig. 2F’, Fig. S1B). A complementary construct lacking only this small region (construct G: −4301 to −1906 / −1598 to +886), shows a reciprocal expression pattern posterior of the furrow with expression restricted to presumptive photoreceptors (Fig. 2G,G’). The 1.2 kb region located at the 5’ end of the *glass* enhancer fragment in combination with the minimal promoter (construct H: −4301 to −3123 / −239 to +886), also restricts GFP expression to cells posterior of the morphogenetic furrow (Fig. 2H). In this case GFP is only expressed in three of the eight presumptive photoreceptors (Fig. 2H’). We identified these as R2, R5, and R8 using defined markers [10] (Fig. S1C). In addition, this part of the *glass* enhancer is required for expression in the ocelli anlage (Fig. 2G,H arrows). Finally, the fragment between the two BamHI sites (construct I: −3123 to −1906 / −239 to +886) drives GFP expression in all presumptive photoreceptors (Fig. 2 I,I’, Fig. S1D), similar to construct D, but with lower expression levels directly after the furrow and increasing GFP levels towards the posterior end.

**Figure 2:**
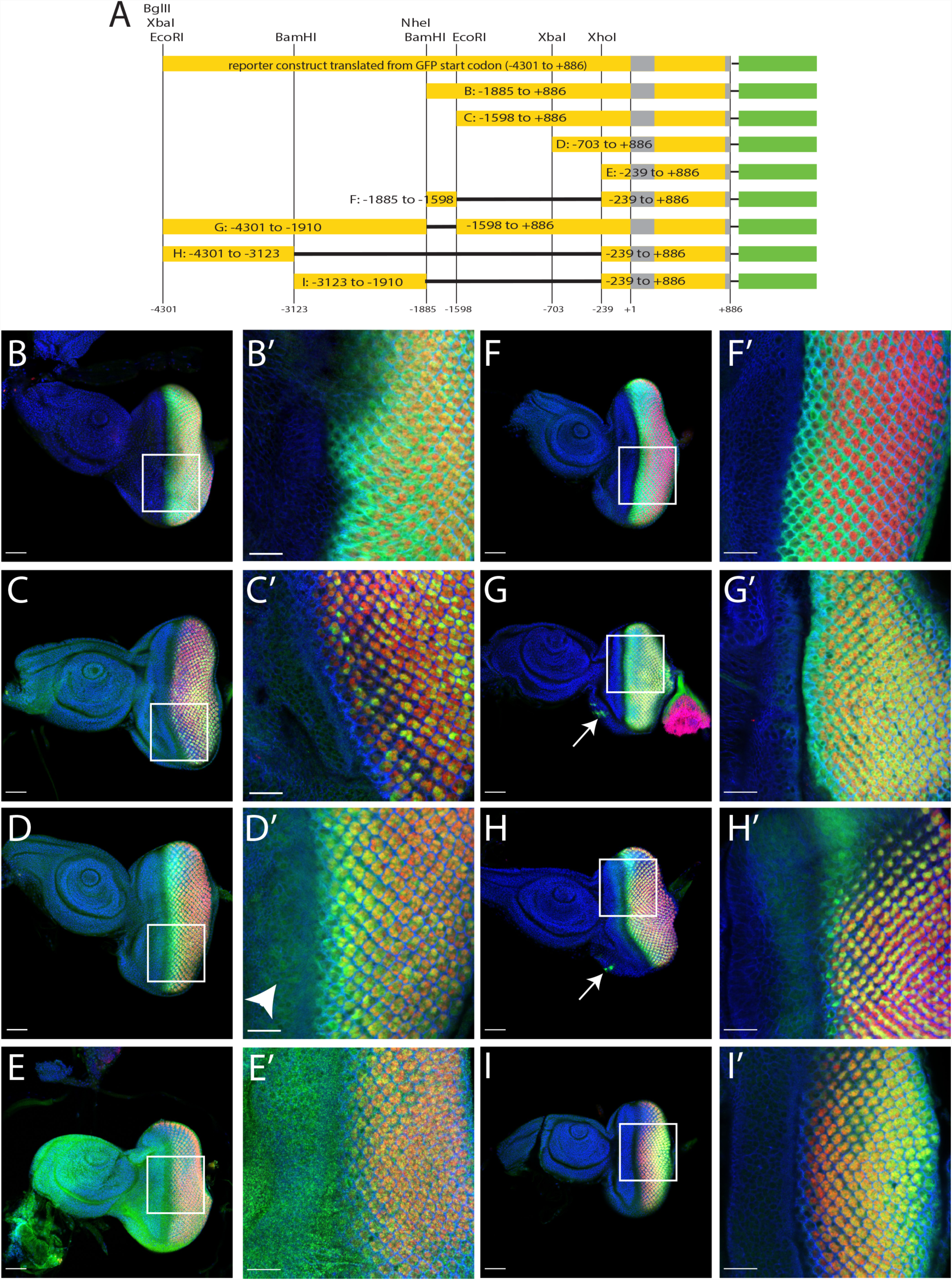
Functional analysis of the *glass* enhancer region. A: enhancer fragments used to drive GFP expression. The construct with the deleted upstream start codons (top) was fragmented using the restriction sites indicated above. The resulting reporter constructs are shown below. B-I: GFP expression patterns of constructs B to I. All discs are oriented with the posterior end to the right. Scale bas: 50µm. Arrows: ocelli anlage. B’–I’: close ups of the regions marked by the rectangles in panel B to I. Scale bars: 20 µm. Arrow head: GFP expression anterior of the morphogenetic furrow. Discs were stained with antibodies against GFP (green), Elav (red), FasIII (blue in panels B’ to I’), and with DAPI (blue in panels B-I)

Taken together 5.2 kb *glass* regulatory region contains a general promoter region (−239 to +886), an ocelli enhancer region (−4301 to −3123, that also drives expression in a subset of photoreceptor precursors, a non-photoreceptor enhancer element (−1886 to −1598), and two general photoreceptor enhancer elements (−3123 to −1906 and −1598 to −239).

### The overlapping upstream smORF is conserved and attenuates Glass misexpression

The upstream start codons in the 5’UTR of *glass* strongly reduced the expression of our original GFP reporter construct, presumably due to interference with GFP translation and production of a 316 amino acid long protein encoded in the 3^rd^ frame of the eGFP sequence used here. In the *glass* transcript, translation from the smORF might also interfere with Glass translation producing a 34 amino acid long peptide overlapping with the Glass coding sequence. Interestingly, the 4 nucleotide sequence preceding the upstream start codon (CAAG) is more similar to the *Drosophila* consensus Kozak sequence (MAAM, whereby M stands for either A or C) [11] than the sequence upstream of the actual Glass start codon (TGTC) (Fig. 3A). Sequence comparison with *glass* genes from other Diptera revealed that upstream start codons are present in all *glass* 5’UTRs of Drosophilidae as well as in *Lucilia, Musca,* and *Glossina,* possibly producing peptides that overlap with the Glass coding sequence (Fig. S2A). Although the length of these peptides differs slightly due to insertion and deletion of nucleotide triplets, the frameshift relative to Glass and the amino acid sequence are conserved within the Drosophilidae, suggesting that the encoded peptide itself might have a conserved function (Fig. S2B). Interestingly, the N-terminal half of the peptide contains mainly basic residues that can provide a nuclear localization signal, as revealed in the GFP reporter construct that was cloned in frame with the upstream start codon (Fig. 1D). The central part of the peptide sequence is more variable and truncated in *D. grimshawi, D. virilis,* and *D. mojavensis,* while it is extended in *D. wilistoni, L. cuprina, M. domestica*, and *G. morsitans* (Fig. S2B). Not surprisingly, conservation is also high in the C-terminal part overlapping the Glass coding sequence. The N-termini of the Glass orthologs of other insects, including mosquitoes, are not conserved, and there are no upstream overlapping open reading frames in these transcripts.

**Figure 3:**
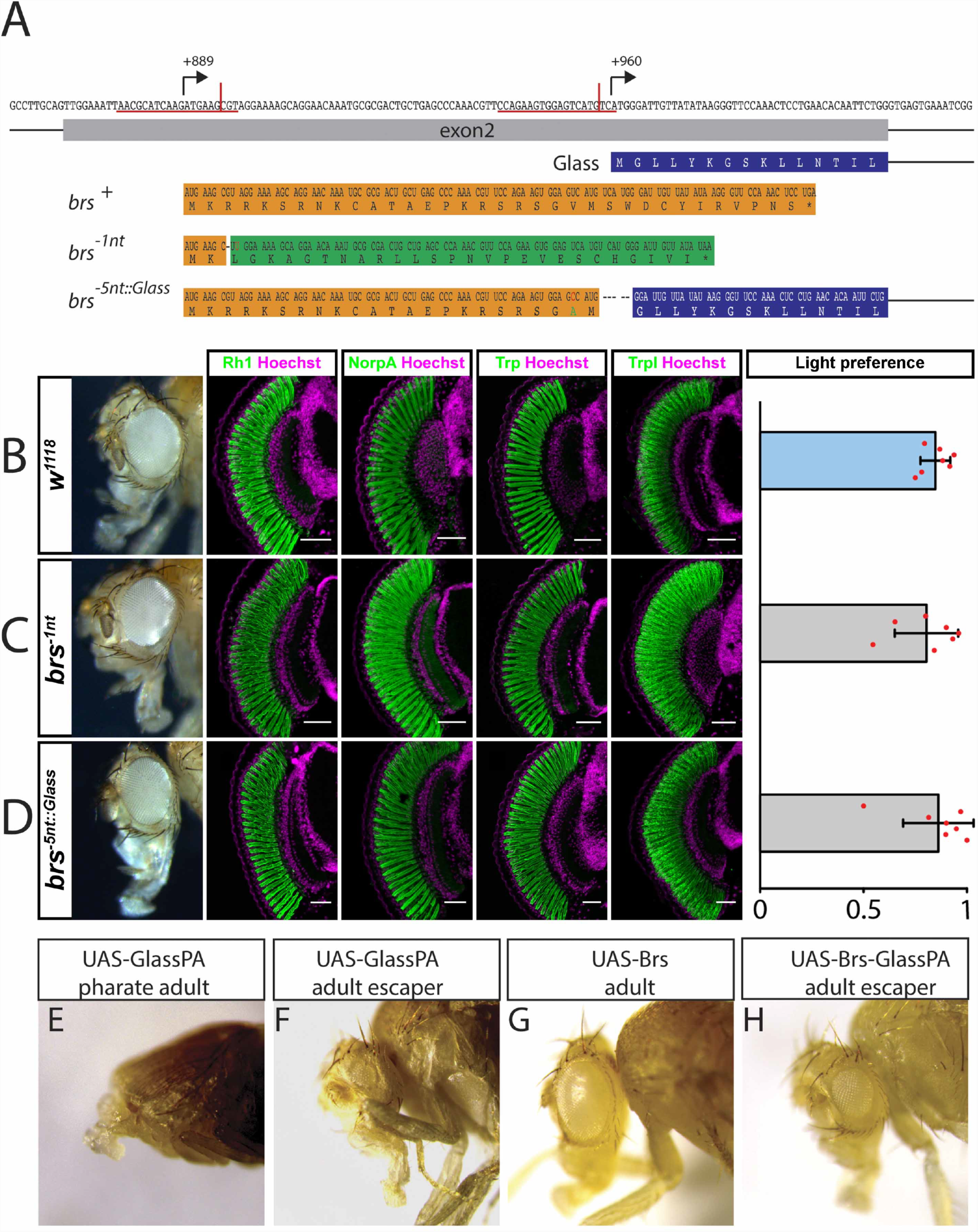
The upstream overlapping open reading frame is not required for eye development or function. A: sequence of *glass* exon 2. The positions of the upstream start codon (+889) and the Glass start codon (+960) are indicated by arrows. CRISPR sites are underlined in red, and the actual positions of the cuts are indicated by a vertical red line. The amino acid sequence of the Glass N-terminus is written in the blue box under the coding sequence. The *brs* nucleotide and amino acid sequence is shown in the orange box. A single nucleotide deletion was introduced in the *brs^-1nt^* allele (followed by an A to T mutation to generate a HindIII restriction site). This will result in a frameshift completely changing the amino acid sequence of the Brs peptide (shown in the green box) with only a minimal change in the nucleotide sequence of exon 2. The resulting peptide still overlaps with the Glass open reading frame. A 5 nucleotide deletion was introduced in the *brs*^*-^5^nt::Glass*^ allele between the second Brs AUG and the second codon of Glass, putting both sequences into the same frame as indicated by the orange and blue underlayed sequences (a T to C mutation was introduced to generate an NcoI site, changing the valine at position 22 of this fusion protein into alanine). B: adult eye, expression of the retinal markers Rh1, NorpA, Trp, and Trpl and ERG of control *w*^*1118*^ flies. C: adult eye, expression of the retinal markers Rh1, NorpA, Trp, and Trpl and ERG of *brs^-1nt^* deletion flies. D: adult eye, expression of the retinal markers Rh1, NorpA, Trp, and Trpl and ERG of *brs*^*-^5^nt::Glass*^ deletion flies. B-D: All antibody stainings are shown in green, DNA was stained with Hoechst (purple). Scale bars represent 40 μm. Flies of all tested genotypes are attracted by light. Two-tailed one sample *t* test followed by the Benjamini Hochberg procedure: For all data sets *n* = 7 experiments. CTRL: *p* = 3.6*10^−8^, *t*_*(6)*_ = 31.09; *brs^-1nt^*: *p* = 1 *10^−5^, t_(6)_ = 13.88; *brs*^*-^5^nt::Glass*^: *p* = 1.1*10^−5^, *t*_*(6)*_ =13.38. The light preference index of all experimental groups is not different from the light preference index of the control group. One-way ANOVA of preference indices: *p* = 0.495, *F-Value*_*(5,36)*_ = 0.895. Data show mean and error bars show standard deviation. Red dots indicate means of individual experiments. E-H: overexpression of UAS constructs with a strong *ey*-Gal4 driver. E: Overexpression of the Glass PA protein leads to severe eye and head defects resulting in pharate lethality. F: the only *ey*-Gal4>*UAS-Glass-PA* fly that eclosed had very small eyes. G: Overexpression of the Brs peptide did not affect eye or head development. H: When expressed together on the same UAS-construct, Brs translation interferes with Glass translation resulting in a higher number of escapers that have small or even normal eyes.

Since the 34 amino acid long peptide is encoded by the *glass* mRNA, it might have a function in eye development. We used the CRISPR/Cas9 technique to introduce small deletions in the peptide coding sequence that will result in a frameshift of the peptide without affecting the Glass coding sequence. We named the resulting smORF alleles “*brainy smurf*” (*brs*) after the smurf with the glasses. We introduced a double strand break 6 nucleotides downstream of the start codon of the peptide and provided a template for repair that contained a single nucleotide change as well as a single nucleotide deletion (*brs^-1nt^*) (Fig. 3A). We crossed the G0 flies to flies that had a deficiency of the *glass* locus and selected offspring that showed a subtle rough eye phenotype over these deficiencies. In addition to the single nucleotide deletion provided by the template, we also found several lines that had small indels in the region of the CRISPR site used (Fig. S3A). We used *w*^*1118*^ flies as controls, since most of our *brs*^*-*^ lines were in a *w*^*-*^ background due to crossing the G0 flies with *w;; Df(3R)Exel6178* and the F1 flies with *w;; Dr e/TM3*. *w*^*1118*^ control flies have big round compound eyes expressing the phototransduction proteins Rhodopsin1 (Rh1), No receptor potential A (NorpA), Transient receptor potential (Trp), and Transient receptor potential-like (Trpl) (Fig. 3B). Adult flies are attracted to light in phototaxis experiments [6]. This is also the case for white eyed mutants (Fig. 3B). *brs^-1nt^* homozygous flies have normal eyes, expressing all tested markters and show light preference comparable to wildtype flies (Fig. 3C). We observed the same phototaxis behaviour and marker gene expression in the randomly generated *brs* mutations (Fig. S3B-F).

The GFP reporter constructs demonstrated that the presence of the upstream start codon can interfere with translation from the actual start codon. To test if this is also the case for the translation of Glass, we introduced a 5 nucleotide deletion at the Glass start codon, putting it into frame with the upstream start codon (*brs^-5nt::Glass^*) (Fig. 3A). In this case, Glass translation starts at the upstream start codon that has the “better” Kozak sequence and fuses the nuclear localization signal encoded by the N-terminus of Brs to the Glass protein. We considered that this could result in higher levels of Glass activity that might interfere with eye development. However, we did not observe any changes in marker gene expression, photoreceptor shape, or light preference (Fig. 3D). Thus, the potential increase of Glass protein either does not interfere with its function, or is compensated by other mechanisms.

To test our hypothesis that the upstream start codon might interfere with Glass translation, we used an over-expression assay. Driving *UAS-glass-PA* expression with a strong *eyeless*-Gal4 enhancer results in lethality of the pharate flies (Fig. 3E, Table1). The flies have severe head defects that prevent them from eclosing with only a small number of escapers (0.4%) (Fig. 3F). Overexpression of a *UAS-brs* construct did not affect viability of the flies or their eye shape (Fig. 3G, Table 1), indicating that the small peptide produced does not interfere with eye development. Co-expression of *UAS-glass-PA* and *UAS-brs* inserted on different chromosomes showed a similar level of lethality as *UAS-glass-PA* alone (0,3% survival rate), indicating that the peptide itself does not interfere with Glass function. However, in a construct that contains the peptide coding sequence upstream and overlapping with the Glass PA coding sequence as in the endogenous transcript, the lethality caused by the over-expression of Glass protein was reduced, resulting in a 20.3% survival rate, where the adult escapers had normal or smaller eyes (Fig. 3H). Therefore, the presence of the Brs peptide itself does not reduce Glass levels. *brs* only interferes with Glass translation, when directly placed as an upstream overlapping open reading frame in the *glass* mRNA.

**Table 1:** Brs interferes with Glass translation. A strong *ey-Gal4/CyO* driver line was crossed with different UAS-constructs and the number of eclosed *Cy^+^* and *CyO* flies was determined. the *Cyo* siblings do not contain the ey-Gal4 driver and therefore were taken as reference for the amount of *Cy^+^* flies expected in each experiment. The ratio of *Cy^+^* over *CyO* flies determines the survival rate of flies expressing the UAS-construct.

| UAS-constructs | # of *Cy<sup>+</sup>* flies | # of *CyO* flies | Total | Survival rate |
| --- | --- | --- | --- | --- |
| *Glass-PA* | 1 | 267 | 268 | 0.4% |
| *brs* | 409 | 408 | 817 | 100.2% |
| *brs-glass-PA* | 134 | 661 | 795 | 20.3% |
| *glass-PA; UAS-brs* | 2 | 702 | 704 | 0.3% |

### The Glass PA protein isoform by itself is sufficient for photoreceptor differentiation and function

*glass* encodes a 604 amino acid protein containing a transcriptional activation domain and a DNA-binding domain that consists of five zinc-fingers of which the three C-terminal zinc-fingers were shown to be necessary and sufficient for DNA binding [7, 12]. According to FlyBase (flybase.org), the *glass* gene encodes three different protein isoforms (Fig 1A). The PA isoform contains a complete set of 5 zinc-fingers providing sequence specific DNA binding to the target genes of Glass [3, 13]. In addition to the Glass PA isoform, two other isoforms are predicted to exist based on expressed sequence tags and sequence conservation [14, 15]. Failure to splice out the last intron of the mRNA transcript, results in the production of a truncated 557 amino acid Glass PB isoform lacking the second half of the fifth zinc-finger. This version of Glass cannot bind specifically to its target sequence *in vitro* [12]. Of the 19 cDNA clones whose sequences are available on FlyBase (flybase.org), seven are covering the last and/or the second-last exon, and all seven still contain the last intron, suggesting that this intron is frequently retained in the transcript. The position of the last intron (intron 4 in *Drosophila*) including the stop codon immediately following the exon-intron junction is only present in the *glass* orthologs of Diptera and Lepidoptera Fig. S4A). In the postman butterfly (*Heliconius melipone*) the stop codon is not located immediately after the exon intron junction but 17 basepairs into the intron. Other arthropods do not have an intron at this position.

An extended 679 amino acid long Glass PC isoform, containing all 5 zinc-fingers followed by additional 75 amino acids, is produced by a readthrough of the Glass PA stop codon. The prediction of this longer isoform is based on sequence conservation 3’ of the regular stop codon [16]. A comparison of the sequence following the Glass stop codon within the higher *Diptera* shows conservation on the amino acid level suggesting that the extended protein is produced by a direct misinterpretation of the stop codon without shifting the reading frame (Fig. S4B). The amino acid sequences of the extended Glass proteins from different higher *Diptera* are highly conserved at their N- and C-termini, but have a central region that is rich in histidine residues of very variable length. Particularly the *Musca* and *Lucilia* Glass PC versions contain a high number of additional amino acids in this central part. The PC sequence is not conserved in other insects including mosquitoes.

To test the requirement and function of the three different Glass isoforms *in vivo*, we introduced specific changes in the endogenous *glass* locus by CRISPR/Cas9-mediated genome editing, eliminating either one or two of the Glass isoforms (Fig. 4A). We used *w*^*1118*^ flies as controls, since our *glass* deletion lines were in a *w*^*-*^ background due to crossing the G0 flies with *w;; Df(3R)Exel6178* and the F1 flies with *w;; Dr e/TM3* (Fig. 4B). Flies expressing only the Glass PA+PC isoforms due to a deletion of the last intron had normal, functional eyes expressing phototransduction proteins like control flies (Fig. 4C). In contrast, a deletion that allowed only the production of the truncated PB isoform phenocopied *glass* amorphic mutations, in which photoreceptors failed to differentiate as revealed by the loss of phototransduction proteins (Fig. 4D) [1]. We also prevented the production of the PC isoform by adding two additional stop codons at the end of the Glass PA sequence. This had no effect on eye shape or on photoreceptor marker gene expression (Fig. 4E). By deleting the last intron and adding stop codons at the end of the Glass PA sequence we generated flies that can only express the PA isoform. These flies also have normal functional eyes expressing all tested markers (Fig. 4F). In addition to marker gene expression, we also measured photoreceptor activity by recording electroretinograms (ERGs). We found that all isoform mutants that had normal eye shape and were expressing phototransduction proteins, showed normal ERG responses [17], while the flies expressing only the Glass PB isoform did not produce any ERG signal in response to light (Fig. 4G). When we tested the light preference of our different Glass isoform mutants, we found that all variations expressing the Glass PA isoform showed light preference comparable to wildtype flies (Fig. 4H). In contrast, the flies expressing only the Glass PB isoform were photoneutral, with a light preference index that is not significantly different from chance, but significantly different from that of control flies and similar to that of *glass* mutants, which fail to detect light [6]. A *glass* mutant phenotype was also observed in flies in which, after CRISPR-induced DNA double strand break, the DNA repair occurred in form of non-homologous end joining, either deleting the exon-intron junction and the stop codon located in the last intron, or introducing a frameshift at the beginning of the last exon (Fig. S5A). Like flies expressing the truncated Glass PB isoform, these flies also have small eyes with a glassy surface. They do not express any of the tested PR makers, have no ERG response and do not show phototaxis behaviour (Fig. S5B-G).

**Figure 4:**
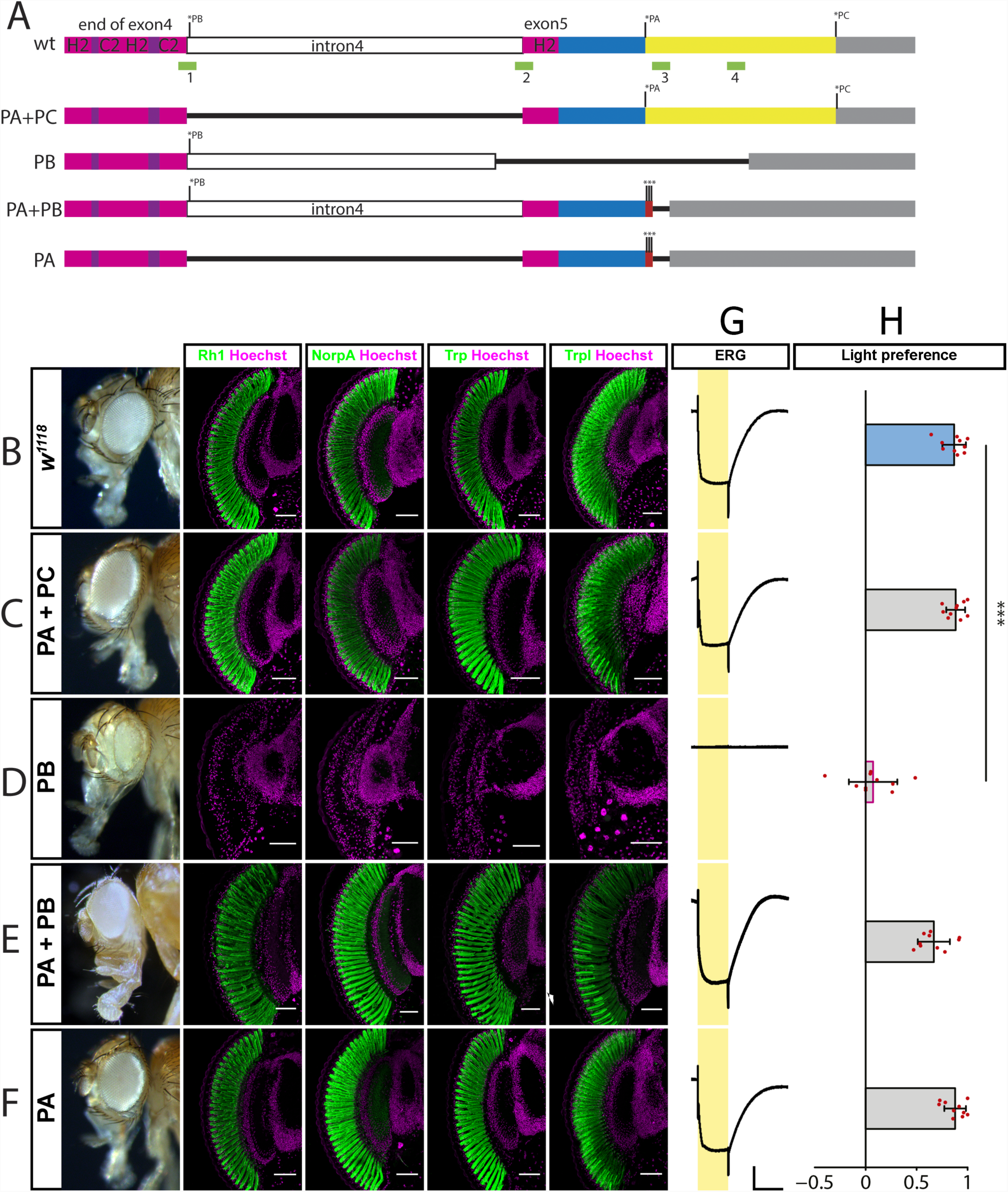
Isoform specific *glass* alleles. A: wildtype and mutated versions of *glass* from the end of exon 4 to the end of the transcript. Stop codons of isoforms PA, PB, and PC are indicated by asterisks. The C2H2-zinc-finger region is shown in purple and magenta. Intron 4 is shown in white. The C-terminus of the PA isoform is shown in blue, that of the PC isoform in yellow. The 3’UTR is grey. The positions of the CRISPR sites used for mutagenesis are shown as green boxes. Deletions are indicated as black lines. The triple stop codon introduced in the PA+PB and the PA alleles are indicated by red boxes and asterisks. B: adult eye and expression of the retinal markers Rh1, NorpA, Trp, and Trpl in control *w*^*1118*^ flies as indicated above. C: adult eye and expression of the retinal markers Rh1, NorpA, Trp, and Trpl in flies expressing the Glass PA +PC isoforms. D: adult eye phenotype and loss of expression of the retinal markers Rh1, NorpA, Trp, and Trpl in flies expressing the Glass PB isoform. E: adult eye and expression of the retinal markers Rh1, NorpA, Trp, and Trpl in flies expressing the Glass PA +PB isoforms. F: adult eye and expression of the retinal markers Rh1, NorpA, Trp, and Trpl in flies expressing the Glass PA isoform. B-F: All tested markers are expressed in the different alleles, except in the mutant only expressing the PB isoform, which has a *glass* mutant eye phenotype. All antibody stainings are shown in green, DNA was stained with Hoechst (magenta). Scale bars represent 40 μm G: ERGs of the different *glass* isoform alleles indicated at the left, scale bars represent 5 mV (vertical) and 5 seconds (horizontal). H: Light preference index of wildtype and flies that are mutant for specific Glass isoforms. Flies expressing only the PB isoform are photoneutral. Two-tailed one sample *t* test followed by the Benjamini Hochberg procedure: For all data sets *n* = 10 experiments. CTRL: *p* = 5.8*10^−09^, *t*_*(9)*_ = 23.82; PA+PC: *p* = 2.1*10^−09^, *t*_*(9)*_ = 30.20; PB: *p* = 0.46, *t*_*(9)*_ = 0.97; PA+PB: *p* = 6.6*10^−07^, *t*_*(9)*_ = 13.43; PA: *p* = 3.8*10^−09^, *t*_*(9)*_ = 26.19. Only flies expressing just the Glass PB isoform show a light preference which is different from the one of control flies. One-way ANOVA of preference indices: For all data sets *n* = 10 experiments, *p*<2*10^−16^, *F-Value*_*(8,81)*_= 44.04. CTRL vs PA+PC: *p* = 1, *t* = −0.15; CTRL vs PB: *p*<0.001, *t* = 8.11; CTRL vs PA+PB: *p* = 0.23, *t* = 2.03; CTRL vs PA: p = 1, *t* = −0.10. Data show mean and error bars show standard deviation. Red dots indicate means of individual experiments. *** = *p*<0.001.

Thus, although conserved, the extended version of Glass is dispensable in photoreceptors. In contrast, the truncated PB version alone cannot fulfil Glass function, while its absence does not interfere with Glass function in the eye.

## DISCUSSION

Changes of the genomic context in a given locus can have a profound influence on gene expression. Addition or deletion of transcription factor binding sites does not only affect when and where a gene is transcribed, but can also determine the expression level. The addition of an exon containing a start codon upstream of the first coding exon can result in an N-terminal extension of the protein if the start codon is in the same reading frame as the coding sequence, or it might interfere with translation if it is in a different reading frame. Insertion of an intron within the coding sequence can result in the production of a truncated protein due to intron retention. Stop codon readtrough can lead to the production of an elongated version of the protein. Such changes can reduce the protein levels or even alter protein function meaning that they are usually quickly removed from the genome. Thus, conservation of such traits over more than the most closely related species indicates that they are neutral or even beneficial. The *Drosophila* transcription factor *glass* combines such features in its transcript, making it a good candidate to investigate these phenomena in the well studied context of photoreceptor development and function. Here we identified an upstream overlapping open reading frame affecting Glass translation. We analysed the role of the different Glass isoforms generated by intron retention and stop codon readthrough. In addition, we dissected the *glass* regulatory sequence and identified several cell- and tissue specific enhancer elements.

### General and cell-specific enhancer elements regulate *glass* expression

The 5.2 kb region upstream of the *glass* start codon had previously been identified as the minimal sequence required for normal Glass expression [8]. The lacZ-reporter construct used in this paper also contained the upstream start codon located in intron 2. Due to the enzymatic activity of the β-galactosidase sufficient signal was produced to detect reporter gene activity posterior of the morphogenetic furrow. However, further truncations of the upstream sequence only yielded transgenic lines with weak or variable expression or lines that did not show expression at all. Similarly, our original GFP-reporter construct showed only very weak expression levels even with the same 5.2 kb enhancer fragment. After removal of the upstream start codon, expression of our GFP-reporter construct was strongly enhanced allowing us to perform a classical enhancer bashing approach to further dissect the upstream region of *glass*. We were able to identify different enhancer regions that conferred reporter gene expression in cells posterior to the morphogenetic furrow. The retinal determination network consisting of several transcription factors, specifies the position of the eye field in many different organisms [18]. Sine oculis, a member of the retinal determination network regulates *glass* expression by directly binding to sites in the enhancer sequence [1]. The three Sine oculis binding sites tested in this paper are located in the BamHI-EcoRI fragment (−1885 to −1598). However, many additional potential Sine oculis binding sites are spread along the entire 5.2 kb upstream sequence (altogether 10 sites with a perfect AGATAC consensus sequence and 22 sites with a more degenerate version YGATAY). Given that all enhancer fragments we tested, showed GFP expression posterior to the morphogenetic furrow, we propose that Sine oculis binds to multiple sites in the 5.2 kb enhancer to activate *glass* expression. In addition to this general reporter gene activation we identified specific enhancer regions driving expression in distinct cell types. For example, the 5’-end of the enhancer that leads to expression in the ocelli anlage and in a subset of the photoreceptors, or the BamHI-EcoRI region that activates expression in non-photoreceptor cells. Thus, other transcription factors binding more specifically to these enhancer elements, might regulate *glass* expression in a cell-type dependent manner.

### The extended Glass PC isoform is dispensable for eye development and function

Stop codon readthrough is relatively abundant in *Drosophila* [19]. Especially genes expressed in the nervous system are putative candidates for this process [20]. Glass has also been listed as a candidate for this protein extension mechanism based on several criteria [16]: Sequence comparison of the amino acids following the regular stop codon shows a higher conservation within higher Diptera than what is found in the 3’UTR of non-readthrough transcripts. The pattern of nucleotide substitutions also suggests that there is no alteration of the reading frame as it might occur in the case of alternative spicing or ribosome hopping. The most frequent stop codon readthrough context (UGAC) [16], is also found at the Glass PA stop codon. Upon readthrough of a UGA stop codon either arginine, cysteine, serine, or tryptophane can be inserted by a near-cognate tRNA at this position [21]. In our isoform deletion experiments we did not test the conditions at which the stop codon is deleted or replaced by another codon (Glass PC and Glass PB+PC) because the function of the resulting protein might be affected by the type of alteration we introduce. Our results from the mutants expressing only the Glass PA isoform suggest, that under laboratory conditions, stop codon readthrough and production of the extended Glass PC version is not required for eye development and photoreceptor function.

### Splicing of intron 4 is required to produce functional Glass protein

The Glass PB isoform alone is not functional. Our deletion mutants that can only express this truncated version as well as our other deletions that affect splicing and result in proteins terminating in intron 4, have a *glass* mutant eye phenotype, that is: they lack the expression of photoreceptor markers, show no photoreceptor activity and are photoneutral [1, 6, 22]. This corroborates previous results that showed that the last three zinc-fingers are essential for sequence specific DNA-binding and that a Glass protein lacking the C-terminal end shows no transcriptional activity [12]. The intron that is retained in the Glass RB transcript, is only found in Diptera and Lepidoptera, suggesting that it originated in the last common ancestor of flies and butterflies. Intron retention can be a means of regulating protein levels since they are usually degraded by nonsense-mediated decay [23]. One of the first examples for cell-type specific intron retention was the *Drosophila* P-element [24]. In germ cells intron 3 is spliced out resulting in functional transposase production. In contrast, in somatic cells intron 3 is retained resulting in a truncated protein that antagonizes the full-length protein. Intron retention can also generate new protein isoforms like the *Drosophila* Noble protein [25]. In addition, intron-retaining mRNA transcripts can remain in the nucleus and be spliced upon requirement providing a source of transcript that could be faster activated than by de novo transcription [26]. Recent RNAseq data suggests that intron 4 is not retained in the *glass* mRNA [4]. The authors found that expression of either the *glass* RA+RC or the *glass* RB transcript from transgenic constructs resulted in production of functional Glass protein, suggesting that in their ectopic expression experiments, intron 4 of the RB transcript was spliced out to produce full-length Glass PA (and PC) protein. Thus, we would consider the *glass* RB transcript as an intermediate stage that has been accumulated during cDNA preparation but that can be further processed to encode functional Glass protein. As we show here, the absence of intron 4 in the *glass* gene allowing only the production of the Glass PA (and PC) protein does not affect eye development and function.

### Brs interferes with Glass translation

According to the scanning model of translation initiation [27], the 40S ribosomal subunit scans the mRNA from the 5’end until it encounters the first AUG codon. Translation will start at this codon, which in the case of *glass* mRNA would mean that only the Brs protein should be produced. However, under certain conditions, translation can also start at a later AUG codon [28]. One of these mechanisms, called leaky scanning, applies for an upstream AUG with a weak context, were the codon with the weak context is skipped by some ribosomes starting translation further downstream. However, this cannot be the case for Glass translation, since there are two AUG codons upstream of the Glass start codon within exon 2 and both have a better Kozak sequence than the actual Glass AUG. Another mechanism would be reinitiation, where after translation of a small upstream open reading frame, the ribosome can move on and re-acquire a Met-tRNA allowing it to reinitiate translation at the next AUG codon. However, since ribosomes cannot backup, the overlapping open reading frame should profoundly inhibit Glass translation [29, 30]. It was shown that overlapping upstream open reading frames; and particularly those that have an optimal AUG context, are efficiently removed from the *Drosophila* genome [31], suggesting that those that can be found and that are even conserved outside of the most closely related species, have been selected due to a specific function. In the case of Glass, we found evidence that translation from the upstream start codon strongly reduces GFP translation and also interferes with Glass translation when overexpressed. However, this suggests that the endogenous Glass protein would be expressed at very low levels. One possible way to overcome this problem, would be by splicing out exon 2 so that the two upstream AUGs and the Glass AUG would be removed from the transcript. In this case translation would start from an AUG codon in exon 3 (amino acid 26 of the predicted full length protein). However, there is no evidence, that exon 2 is spliced out of the transcript to produce a truncated Glass version. Another hypothesis would be that Glass translation starts by reinitiation at the AUG codon in exon 3 after translation of Brs. In fact, the Glass proteins of *Anopheles darlingi* and of *Culex quinquefasciatus* are predicted to start at this position (with a conserved motif: MYISC), while *Anopheles gambiae* Glass is predicted to start in an exon located further upstream of the start codons of the other two mosquito species, but with an N-terminal sequence that is not related to that found in *Drosophila* and other higher Diptera. Also, other insects’ Glass proteins start further downstream than in the Diptera. It could be possible that for most insects the actual Glass translation start has not yet been identified due to higher sequence divergence at the N-termini. This would suggest, that the first 25 amino acids mostly encoded by exon 2 of the *Drosophila* transcripts are dispensable for Glass function, or that they are only required in higher Diptera. In addition to regulating Glass protein levels by directly interfering with translation efficiency, the Brs peptide could have other functions in the developing eye, where it is expressed along with Glass. Small peptides can have important roles such as hormones, pheromones, transcriptional regulators, antibacterial peptides, etc. [32]. However, we could neither identify such a role for Brs by mutating it, nor by overexpressing it, suggesting that its main role is the regulation of Glass translation.

In summary, our results suggest that the removal of intron 4, which was added in the common ancestor of flies and butterflies, is essential for the production of a functional Glass protein. Stop codon readthrough resulting in an extention of the Glass protein that is conserved in higher Diptera seems to be dispensable for Glass function in photoreceptor development. The addition of an exon containing several AUGs upstream of the Glass start codon found in mosquitoes, can interfere with Glass translation. Nevertheless, conservation of the upstream start codon and sequence conservation of the Brs peptide suggest that higher Diptera have found a way to overcome this interference and that Brs might even have adopted a beneficial function.

## MATERIAL AND METHODS

### Fly strains

Flies were reared at 25°C on a cornmeal medium containing agar, fructose, molasses, and yeast. Strains for site directed integration (25709, 25710), *w*^*1118*^ mutants (3605), deficiency lines (4431, 7657) [33], balancer lines (36305, 8379), and *nos-Cas9* expressing flies (54591) [34] were obtained from the Bloomington stock center. *ey-Gal4* expressing flies were a kind gift from R. Stocker.

### Transgenic constructs

All oligos used for cloning and all sequencing reactions were purchased from microsynth. The following primer sequences show the annealing sequence part in capitals and any additional sequence in small letters. Restriction sites are underlined.

A 5257 basepair long PCR fragment was amplified from genomic DNA of CantonS flies using primers “glass −4301 Asc fw” (5′-<u>ggcgcgcc</u>TAACCCGATACAAATGGAGAGG-3′) and glass 5’UTR Not re” (5′-<u>gcggccgc</u>GACATGACTCCACTTCTGGAAC-3′). The fragment was inserted into pCR-Blunt II-TOPO vector (Invitrogen). From there it was excised using the restriction enzymes AscI and NotI and cloned into a GFP reporter vector (pDVattBR, kind gift from Jens Rister) to generate the basic *glass*-GFP reporter construct (Fig. 1A, B’). The plasmid was injected into y, w; attP2 embryos to produce transgenic flies (Genetic services Inc.).

To delete the two upstream start codons, a 1483 bp PCR product was amplified from the original glass-GFP reporter plasmid using primers “glass −597 fw” (5’-TAAAAACTACTGAAAACTGCTGCCGATG-3’) and “glass exon2 noAUG Pme re” (5’-gc<u>gtttaaac</u>GATGCGTTAATTTCCAACTGCAAGGC-3’), TOPO cloned into pCRII, sequenced, digested XhoI-PmeI, and transferred into the original plasmid also cut with XhoI-PmeI, thereby removing the 104 basepairs encoding the N-terminus of Brs, and part of the multiple cloning site of the vector (Fig. 1C’).

To put the GFP coding sequence in frame with the upstream start codon, the GFP coding sequence was amplified by PCR using the primers “GFP noStart Not fw” (5’-tc<u>gcggccgc</u>gGGTGAGCAAGGGCGAGG-3’) putting a NotI site in front of GFP (starting with the 3^rd^ nucleotide of GFP) and “GFP down Fse re” (5’-GATTATGA<u>TCTAGA</u>GTCGC<u>GGCCG</u>-3’) covering an FseI and an XbaI site in the plasmid. The PCR product was cloned NotI-XbaI into pBluescript, sequenced, and transferred NotI-FseI into the original glass-GFP reporter plasmid deleting most of the multiple cloning site and the GFP start codon (Fig. 1D’).

For the enhancer analysis, the construct lacking the upstream start codons was digested with different combinations of restriction enzymes and religated. For construct B the plasmid was digested with BglII cutting in the multiple cloning site at the 5’-end of the enhancer and with BamHI cutting at position −3123. A second BamHI site at position −1885 was not in the database sequence but is present in the fragment amplified from the CantonS flies. Therefore, religation of the plasmid after BglII-BamHI digestion (the two enzymes producing compatible sticky ends) resulted in an enhancer fragment ranging from position −1885 to +886. For construct C the plasmid was digested with EcoRI cutting at the multiple cloning site at the 5’-end of the enhancer and at positions −1598 and −2040 in the enhancer. Religation resulted in an enhancer fragment ranging from position −1598 to +886. For construct D the plasmid was digested with XbaI cutting in the multiple cloning site at the 5’-end of the enhancer and at position −703. Religation resulted in an enhancer fragment ranging from −703 to +886. For construct E, construct C was digested with XbaI cutting in the multiple cloning site at the 5’-end and at position −703 in the enhancer as well as with XhoI cutting at position −239 in the enhancer. The two ends were filled using Klenow polymerase and religated resulting in an enhancer fragment ranging from position - 239 to +886. For construct F, construct B was digested with EcoRI cutting at position −1598 and XhoI cutting at position −239. The two ends were filled using Klenow polymerase and religated resulting in an enhancer fragment ranging from position - 1885 to −1598 fused to the minimal promoter fragment from position −239 to +886. For construct G, the original −4301 to +886 plasmid was digested with NheI cutting at positions −1910, −1903, and −682, the site at position −2179 is missing in our enhancer fragment amplified from CantonS flies. In another reaction the original plasmid was digested with EcoRI cutting at positions −1598 and −2040. Both reactions were filled using Klenow polymerase, digested with BglII, and then the BglII-NheI_filled_ fragment was ligated into the BglII-EcoRI_filled_ fragment fusing the enhancer region from −4301 to −1910 to the region from −1598 to +886. For construct H, the original plasmid was digested BamHI-XhoI. The two ends were filled using Klenow polymerase and religated to fuse the fragment from −4301 to −3123 to the fragment from −239 to +886. For construct I, the original plasmid was digested with NheI and in an independent reaction with XhoI. Both digestion reactions were filled with Klenow polymerase. The NheI_filled_ reaction was further digested with BamHI and the XhoI_filled_ reaction was further digested with BglII. Then the BamHI-NheI_filled_ fragment was ligated into the BglII-XhoI_filled_ plasmid resulting in a fusion of an enhancer fragment ranging from - 3123 to −1910 to the minimal promoter ranging from −239 to +886.

All constructs were injected into *nos-ΦC31;; attP2* flies for site directed integration. The G0 flies were crossed individually to *w*^*1118*^ flies to screen for *w*^*+*^ offspring. *w*^*+*^ F1 flies were crossed individually to 3^rd^ chromosome balancer flies (*w;; Dr e/TM3*) and their balanced offspring was crossed inter se to produce stable lines.

For the UAS constructs we used the *glass* cDNA plasmid GH20219 as starting point. This cDNA still contains intron 4 due to incomplete splicing resulting in the RB transcript isoform. To remove the intron, two PCR reactions were set up. One with primers “glass 5’UTR BamHI fw” (5’-ga<u>ggatCC</u>TCGCCAAAAGTCGCTTCTTG-3’) and “glass exon4 re” (5’-ccccgactgcgaaaatCTGAGCAGGCAGAGCTTGCAC-3’) resulting in a fragment ranging from the 5’-end of the 5’UTR to the end of exon 4, with the sequence given in small letters of the reverse primer overlapping with the beginning of exon 5. The other PCR reaction was done with primers “glass exon5 fw” (5’-gctctgcctgctCAGATTTTCGCAGTCGGGGAACTTG-3’) and “gl Stop Xho re” (5’-gg<u>ctcgaG</u>TCATGTGAGCAGGCTGTTGCC-3’), resulting in a fragment ranging from the beginning of exon 5 to the PA stop codon, with the sequence given in small letters of the forward primer overlapping with the end of exon 4. Both PCR products were mixed together to provide the template for another PCR reaction with primers “gl 5’UTR BamHI fw” and “gl Stop Xho re”. The resulting PCR product ranging from the 5’UTR to the PA stop codon without intron 4 was digested with BamHI-XhoI and cloned into pBluescript. After sequencing, different fragments were PCR amplified. The Glass PA coding sequence was amplified with primers “gl Start+Kozak attB1 fw” (5’-ggggacaagtttgtacaaaaaagcaggcttcaaCATGGGATTGTTATATAAGGGTTCCAAACT-3’) and “gl Stop attB2 re” (5’-ggggaccactttgtacaagaaagctgggtcgTCATGTGAGCAGGCTGTTGCC-3’). The *brs* sequence was amplified with primers “glass+Smurf attB1 fw” (5’-ggggacaagtttgtacaaaaaagcaggcttcCGCATCAAGATGAAGCGTAGGAAAAGC-3’) and “glass Smurf Stop attB2 re” (5’-ggggaccactttgtacaagaaagctgggtcTCAGGAGTTTGGAACCCTTATATAACAATCCC-3’). The *brs-glass-PA* sequence was amplified with primers “glass+Smurf attB1 fw” and “gl Stop attB2 re”. The primer sequence in small letters are the attB parts used for gateway cloning. The PCR products were gateway cloned into pENTRY201, sequences, and transferred into the vector *pUASg.attB* for injection (Genetic Services Inc.). *UAS-glass-P*A and *UAS-brs-Glass-PA* were injected into *nos-ΦC31; attP40* flies, while the *UAS-brs* plasmid was injected into *nos-ΦC31;; attP2* flies. After balancing the transgenic flies, *UAS-glass-RA*^*attP40*^ and *UAS-brs*^*attP2*^ were combined in a single line: *w; UAS-GlassRA*^*attP40*^; *UAS-Brs*^*attP2*^. The different UAS-construct bearing flies were crossed to *ey-Gal4/CyO* flies, and the number of offspring with *Cy*^*+*^ versus the number of offspring with *CyO* wings was determined. For calculation of the survival rate the number of *Cy*^*+*^ flies was divided by the number of *CyO* flies (Table 1).

### CRISPR

For the alterations of the endogenous glass locus by CRISPR/Cas9 genome editing, we assembled the different templates in pBluescript. For the Glass PA+PC variant, we needed to remove intron 4 from the genomic DNA without changing the sequence at the Glass PA stop codon. Since there were no useful restriction sites between the intron 4 / exon 5 junction and the Glass PA stop codon, we decided to introduce an NdeI site in this sequence by altering a single nucleotide in the third position of the codon for the first histidine residue of the last zinc-finger (histidine 567 of Glass PA: CAC to CAT). We PCR amplified a 932 bp fragment from the genomic DNA of *nos-Cas9* flies using primers “glass ex5 R1 Nde fw” (5’-ga<u>gaattcatATG</u>CGCGTCCACGGCAAC-3’) and “glass 3’UTR re” (5’-GATCAAAGCACCTGTCTTACATCTACGTCTAG-3’), and a 1529 bp fragment from the intronless glass version assembled in pBluescript for generation of the *UAS-glass-PA* construct using primers “glass ex4 R1 fw” (5’-cg<u>gaattC</u>AAGAGTGCGCCGCTTCC-3’) and “glass ex5 Nde re” (5’-CG<u>CATaTG</u>CCGATTCAAGTTCCCCGAC-3’). Both PCR products were combined in pBluescript vector by digesting the one covering the C-terminus from the NdeI site introduced in exon 5 to an endogenous HindIII site in the 5’UTR with NdeI-HindIII, and the one covering exon 4 and part of exon 5 with EcoRI-NdeI (due to an endogenous NdeI site in the middle of exon4 this part was cloned in two steps). The sequence of the resulting fragment ranging from the beginning of exon 4 to the 5’UTR and lacking intron 4 was confirmed.

For the Glass PB variant, we introduced a deletion ranging from the end of intron 4 to the middle of the sequence added in the extended Glass PC protein version (Fig. 4A). We amplified two PCR products from the genomic DNA of *nos-Cas9* flies. A 1866 bp fragment spanning exon 4 and most of intron 4 was amplified with primers “glass ex4 R1 fw” and “glass int4 Xho re” (5’-AA<u>CTCGAg</u>GTATAACGTTCCAGGACTGCTC-3’),and a 1287 bp fragment ranging from the middle of the Glass PC encoding sequence to a place located around 5000 bp downstream of the *glass* gene was amplified with primers “glass 3’UTR Xho fw” (5’-aa<u>ctcgAG</u>CATCGGCGATTATACTCCACC-3’) and “glass down Kpn re” (5’-ag<u>ggtACC</u>TTTATGGTGGCCTCCCAGG-3’). Both fragments were subcloned, sequenced and combined in pBluescript by digesting the one covering exon 4 and intron 4 with EcoRI-XhoI, and the one covering part of the PC coding region, the 3’UTR and donwnstream genomic sequence with XhoI-Acc65I.

For the Glass PA+PB variant, we introduced two additional stop codons at the end of the Glass PA encoding part and deleted 29 of the nucleotides following the stop codon. We PCR amplified two PCR products from the genomic DNA of *nos-Cas9* flies. A 2034 bp fragment spanning exon 4, intron 4, and the PA encoding part of exon 5 was amplified with primers “glass ex4 fw” and “glass RA **3xStop** Xho re” (5’-ct<u>ctcgag</u>**ctattaTCA**TGTGAGCAGGCTGTTGCCAC-3’). A 1365 bp fragment spanning most of the Glass RC specific sequence, the 3’UTR and around 500 nucleotides of downstream genomic sequence was amplified using primers “glass RC Xho fw” (5’-tt<u>ctcgAG</u>CATTACCACCCCCCGC-3’) and “glass down Kpn re”. Both fragments were subcloned, sequenced, and combined in pBluescript by digesting the one covering exon 4, intron 4, and the Glass PA encoding part plus stop codons with EcoRI-XhoI, and the one covering the Glass PC region, 3’UTR, and downstream genomic sequence XhoI-Acc65I.

For the Glass PA variant, we performed a PCR amplification of exon 4 and the 5’-end of exon 5 with the same first primer pair as for the Glass PA+PB template, but using the intronless glass version assembled in pBluescript for generation of the *UAS-glass-PA* construct as a template. We subcloned this 1638 bp fragment EcoRI-XhoI, sequenced it, and combined it with the 1365 bp fragment spanning most of the Glass RC specific sequence, the 3’UTR and around 500 nucleotides of downstream genomic sequence.

For expression of the CRISPR guideRNAs, sense and antisense oligos with overhangs fitting the sticky ends of the BbsI digested vector were annealed and ligated into pU6-BbsI-chiRNA plasmid (a gift from Melissa Harrison & Kate O’Connor-Giles & Jill Wildonger, Addgene plasmid # 45946 [35]). Site 1 (ctgctcaggtgagtccg/gga) is located at the junction between exon 4 and intron 4. Site 2 (gtccacagattttcgca/gtc) is located at the junction between intron 4 and exon 5. Site 3 (agg/agtgcaggaggtttcca) guides Cas9 to cut 12 bp downstream of the Glass PA stop codon. Site 4 (aca/tgggtaactacgactac) is located in the middle of the Glass PC encoding region (Fig. 4A). CRISPR sites were selected based on their position in the glass genomic sequence using a CRISPR site prediction program (http://tools.flycrispr.molbio.wisc.edu/targetFinder/ [36]).

The templates for the different Glass isoform variants were co-injected with sgRNA expression plasmids into embryos of *nos-Cas9* flies. The Glass PA+PC variant was co-injected with the sgRNA plasmids for sites 1 and 2. The template for the Glass PB variant was co-injected with sgRNA plasmids for sites 2 and 4. The template for the Glass PA+PB variant was co-injected with the sgRNA plasmid for site 3. The template for the Glass PA variant was co-injected with the sgRNA plasmids for sites 1 and 3. The resulting G0 flies were crossed individually with deficiency lines uncovering the *glass* locus. Some of the offspring resulting from Cas9 cutting at CRISPR site 1 (and 3) showed a *glass* mutant phenotype due to CRISPR induced non homologous end joining (Fig. S5). Irrespective of the eye phenotype the offspring was crossed individually with third chromosome balancer flies (*w;; Dr, e/TM3*) and analyzed by PCR for the introduced deletions and sequence alterations. The *glass* genes of those lines that showed changes in the PCR analyses, were sequenced to confirm the introduced changes and identify other alterations that resulted in *glass* mutant phenotypes.

For the deletions in the *brs* sequence, template sequences were assembled in pBluescript. For the 1nt deletion two PCR products were amplified. A 2759 bp fragment ranging from position −1863 in the glass enhancer region to position +896 in the *brs* coding sequence was amplified using primers “glass −2700 Kpn fw” (5’-tg<u>ggtacC</u>GGCAGCAGAGACAGGCTC-3’) and “smurf −1nt H3 re” (5’-cc<u>aaGCTT</u>CATCTTGATGCGTTAATTTCCAACTGC-3’). The resulting PCR product was cloned Acc65I-HindIII into pBluescript and sequenced. A 2493 bp fragment ranging from position +899 in the *brs* coding sequence to position +3392 at the end of exon 4 was amplified from *nos-Cas9* genomic DNA using primers “smurf −1nt H3 fw” (5’-ga<u>aagctt</u>GGAAAAGCAGGAACAAATGCGCG-3’) and “glass exon4 Pst re” (5’-ct<u>ctGCAG</u>GCAGAGCTTGCACTGG-3’). The resulting PCR product was cloned HindIII-PstI into pBluescirpt, sequenced, and combined with the 5’-fragment. For the 5 nt deletion fusing Brs with Glass, A 2821 bp fragment ranging from position −1863 to +953 in the *brs* coding sequence just before the second AUG codon was amplified from *nos-Cas9* genomic DNA using primers “glass −2700 Kpn fw” and “smurf −5nt BamH Nco re” (5’-cc<u>ggatccCATGg</u>CTCCACTTCTGGAACGTTTGGGC-3’). The resulting PCR product was cloned Acc65I-BamHI into pBluescript and sequenced. A 1632 bp fragment ranging from position +959 just before the Glass start codon to position +2591 in exon 4 was amplified from *nos-Cas9* genomic DNA using primers “smurf −5nt Nco fw” (5’-ct<u>cCATGG</u>GATTGTTATATAAGGGTTCCAAACTCCTG-3’) and “glass ex4 Not re” (5’-aagcggccgcatggtgcatggtcatgttcatgc-3’), cloned NcoI_filled_-NotI into pBluescript BamHI_filled_-NotI, sequenced, and combined KpnI-NcoI with the 5’-fragment. For expression of the CRISPR guideRNAs, sense and antisense oligos with overhangs fitting the sticky ends of the BbsI digested vector were annealed and ligated into the BbsI digested pCDF4-U6:1_U6:3tandemgRNAs plasmid (gift from Simon Bullock, addgene plasmid # 49411). The site for the −1nt deletion (aacgcatcaagatgaag/cgt) is located at the upstream start codon, the site for the −5nt deletion (ccagaagtggagtcatg/tca) is located at the Glass start codon (Fig. 3A). The templates for the *brs* mutations were co-injected with the corresponding sgRNA expression plasmid into embryos of *nos-Cas9* flies. The resulting G0 flies were crossed individually with deficiency lines uncovering the glass locus. Some of the F1 flies had a very subtle rough eye phenotype over the glass deficiency – but also over the TM6b balancer. The F1 flies were crossed individually with 3^rd^ chromosome balancer flies to establish stocks, and tested by PCR and restriction digest with either HindIII or NcoI for presence of the introduced changes. Genomic DNA of the *glass* locus from homozygous candidates was PCR amplified and sent for sequencing. Additional sequence changes due to non-homologous end joining were also identified (Fig. S3).

### Light Preference Assay

The light preference assays were prepared and performed under red light conditions during the subjective day. *Drosophila melanogaster* adults of both sexes were used. Given that light perception in *Drosophila* is affected by age [6], for consistency, we used <1 day old flies in all our experiments. Without anaesthesia 20 – 40 flies were taken from food vials and loaded into the elevator chamber of a T-maze. The elevator chamber was descended and flies were allowed to move freely between the elevator chamber and two plastic tubes for 2 min. A white LED was placed at one end of one testing tube. Thus, only one testing tube was illuminated. After 2 min the elevator chamber was ascended and flies were not able to move between testing tubes and elevator chamber. We determined the number of flies in the illuminated testing tube (L) and the number of flies in the dark testing tube (D) as well as the number of flies in the elevator chamber (E). We calculated a preference index as follows:

Light intensity was measured from the distance of the elevator chamber to the LED. The light intensity was 1338 µW/cm^2^ with a first maximum intensity peak of 16.6 µW/cm^2^/nm at 443 nm with half-widths of around 11 nm and a second maximum intensity peak of 6.8 µW/cm^2^/nm at 545 nm with half-widths of around 62 nm.

### Electroretinogram

ERG recordings were obtained as previously described [37]. Briefly, we mounted living flies inside a pipette tip, leaving their heads outside, and immobilised them with a mixture of bee wax and colophony 3:1, which worked as a glue. We placed the flies inside a dark chamber and applied two electrodes: a ground electrode was positioned inside the head of the fly, and a recording electrode was introduced into the retina. In our stimulation protocol, we illuminated the compound eye with orange light for 5 seconds to transform all metarhodopsin to rhodopsin, switched off the light for 10 seconds, and illuminated again a second time with orange light for another 5 seconds. We recorded the response of PRs to the second stimulus.

### Immunohistochemistry

Eye imaginal discs were dissected from third instar larvae and fixed in 3.7% formaldehyde disolved in phosphate buffer (PB) for 20 minutes.

For cryosections, we dissected the heads of the flies and fixed them for 20 minutes with 3.7% formaldehyde disolved PB, as previously described [1]. We washed these samples by using phosphate buffer with Triton 0.3% (PBT) and incubated them overnight in cryoprotected solution (25% sucrose in PB). Then, we embedded the heads in OCT, froze them, and took 14 μm sections by using a cryostat.

Both eye imaginal discs and cryosections were washed with PBT, and incubated sequentially in primary and secondary antibodies (each antibody incubation step was performed overnight, washing with PBT between and in the end of these steps). We used Vectashield as a mounting medium.

As primary antibodies, we used rabbit anti-GFP (1:1000, Molecular probes, A-6455), and guinea pig anti-Sens (1:800, courtesy of H. Bellen [38]). To stain against phototransduction proteins we used antibodies generated in C. Zuker’s lab: anti-NorpA (1:100), rabbit anti-Trpl (1:100), both of which were kindly provided by N. Colley. The following antibodies were obtained from Developmental Studies Hybridoma Bank (DSHB) at The University of Iowa: mouse anti-Rh1 (1:20, 4C5), mouse anti-Trp (1:20, MAb83F6), and rat anti-Elav (1:30, No. 7E8A10)

Secondary antibodies were obtained from Molecular probes, and we used the conjugated with the following Alexa fluor proteins: 488, 568, and 647. In addition, we also used Hoechst 33258 (1:100, Sigma, No. 94403)

## ACKNOWLEDGEMENTS

We thank J. Rister and S.Bullock for plasmids; M.Harrison, K. O’Connor-Giles, and J.Wildonger for the pU6-BbsI-gRNA plasmid and the CRISPR target finder website; R. Stocker for the *ey-Gal4* flies; and N. Colley, H. Bellen, and the DSHB for antibodies. We thank J. E. Treisman for helpful comments and discussions. We are also grateful to our colleagues from the Sprecher and Huber labs for valuable discussions, particularly thanks to B. Egger, T. Smylla and O. Voolstra. This work was supported by the Swiss National Science Foundation (grant number 31003A_149499), the Novartis Foundation for Biomedical Resarch (grant number 18A017) and the University of Fribourg to S.G.S.

---

**Figure S1: *glass*-GFP reporter gene expression patterns**. GFP (green), Senseless (Sens, red), and Elav (blue) expression in the eye region of larval imaginal discs of transgenic flies expressing construct C (−1598 to +886) (A), construct F (−1885 to −1598 / −239 to +886) (B), construct H (−4301 to −3123 / −239 to +886) (C), or construct I (−3123 to −1906 / −239 to +886) (D) (compare to Fig. 2A for the individual constructs). Scale barr: 40 µm A’ to D’: magnification of areas in panels A to D. Scale barr: 5 µm. A’’ to D’’: GFP channel alone.

**Figure S2: sequence conservation upstream of the *glass* start codon.** A: nucleotide sequence of glass exon 2 of different higher *Diptera*. The position of the glass start codon is outlined in green. The position of the upstream start codon is outlined in blue. The black vertical lines show the tripplets following the upstream start codon that result in the amino acid sequences shown in B. B: Alignment of the amino acid sequence resulting from translation beginning at the start codon upstream of Glass from different higher Diptera. *Drosophila melanogaster (D mel), Drosophila sechellia (D sec), Drosophila simulans (D sim), Drosophila erecta (D ere), Drosophila yakuba (D yak), Drosophila takahashii (D tak), Drosophila eugracilis (D eug), Drosophila rhopaloa (D rho), Drosophila ficusphila (D fic), Drosophila elegans (D ele), Drosophila ananassae (D ana), Drosophila persimilis (D per), Drosophila pseudoobscura (D pse), Drosophila grimshawi (D gri), Drosophila virilis (D vir), Drosophila mojavensis (D moj), Drosophila willistoni (D wil), Lucilia cuprina (L cup), Musca domestica (M dom), Glossina morsitans (G mor).*

**Figure S3: *brs* alleles generated by small CRISPR deletions in the *glass* 5’UTR.** A: sequence alignment of the different *brs* alleles. The *glass* start codon is outlined in green, the upstream start codon in blue. The second AUG codon in the same frame as the upstream start codon is outlined in orange. Stop codons relative to the upstream start codon(s) are outlined in red. B: Adult eye phenotype and expression of the retinal markers Rh1, NorpA, Trp, and Trpl of *brs*^*-^2^nt*^ flies. C: Adult eye phenotype and expression of the retinal markers Rh1, NorpA, Trp, and Trpl of *brs*^*-^4^nt*^ flies. D: Adult eye phenotype and expression of the retinal markers Rh1, NorpA, Trp, and Trpl of *brs*^*-^5^nt*^ flies. B-D: All antibody stainings are shown in green, counterstaining of DNA with Hoechst (magenta). All tested photoreceptors are expressed in the *brs* frameshift alleles. Scale barrs: 40 µm E: ERGs; scale bars: 5 mV (vertical) and 5 seconds (horizontal). F: Flies of all tested genotypes are attracted by light. Two-tailed one sample *t* test followed by the Benjamini Hochberg procedure: For all data sets n = 7 experiments. *Brs-2nt*: *p* = 3.6*10^−8^, *t*_*(6)*_ = 47.23; *Brs-4nt*: *p* = 8.9*10^−6^, *t*_*(6)*_ = 14.81; *Brs-5nt*: *p* = 2*10^−6^, *t*_*(6)*_ = 19.97. The light preference index of all experimental groups is not different from the light preference index of the control group. One-way ANOVA of preference indices: *p* = 0.495, *F-Value*_*(5,36)*_ = 0.895. Data show mean and error bars show standard deviation. Red dots indicate means of individual experiments.

**Figure S4: sequence conservation of intron 4 and Glass PC.** A: nucleotide sequence of the transition between exon 4 and intron 4 and between intron 4 and exon 5 of *Drosophila melanogaster*. Other Diptera and Lepidoptera also contain this intron followed by a stop codon black box) immediately after the exon intron junction (black line), except in *Heliconius*, where the stop codon is located 17 pb downstream of the exon intron junction. Although the intron is absent in other insect species, the amino acid sequence flanking the intron and forming part of the Glass zinc-finger is highly conserved as indicated by the translation below the alignment. B: amino acid alignment of the Glass C-termini of different higher Diptera. The position of the Glass PA stop codon is marked by an asterisk (arrow). The amino acid sequence directly following the end of the PA isoform is highly conserved. There is also high sequence conservation at the C-terminus of the PC isoform. The central region, which is rich in histidine residues, is more variable. *Drosophila melanogaster (D mel), Drosophila simulans (D sim), Drosophila sechellia (D sec), Drosophila erecta (D ere), Drosophila yakuba (D yak), Drosophila ananassae (D ana), Drosophila pseudoobscura (D pse), Drosophila persimilis (D per), Drosophila willistoni (D wil), Drosophila virilis (D vir), Drosophila mojavensis (D moj), Drosophila grimshawi (D gri), Musca domestica (M dom), Glossina morsitans (G mor), Lucilia cuprina (L cup), Anopheles darlingi (A dar), Anopheles gambiae (A gam), Culex qquinquefasciatus (C qui), Culex pipiens (C pip), Danaus plexipus (D ple), Heliconus melpomene (H mel), Apis melifera (A mel), Bombus impatiens (B imp), Nasonia vitripennis (N vit), Tribolium castaneum (T cas), Pediculus humanus (P hum), Acyrthosiphon pisum (A pim), Ixodes scapularis (I sca), Drosophila ficusphila (D fic), Drosophila eugracilis (D eug), Drosophila biarmipes (D bia), Drosophila takahashii (D tak), Drosophila elegans (D ele), Drosophila bipectinata (D bip), Drosophila kikkawai (D kik).*

**Figure S5: Additional *glass* alleles generated by non-homologous end joining.** A: wildtype and mutated versions of *glass* from the end of exon 4 to the end of the transcript. Stop codons of isoforms PA, PB, and PC are indicated by asterisks. The C2H2-zinc-finger region is shown in magenta. Intron 4 is shown in white. The C-terminus of the PA isoform is shown in blue, that of the PC isoform in yellow. The 3’UTR is grey. The sequences at the exon intron and intron exon junctions are given as letters with a black vertical line depicting the position of the junction. The stop codon at the beginning of exon 4 is highlighted in blue. The positions of the CRISPR sites used for mutagenesis are underlined in green. Deletions are indicated as black lines in the schemes, and as red letters in the sequences. Due to the deletions of the exon intron junction and the stop codons, additional amino acids are added to the Glass PB sequence until they reach the next stop codon in intron 4 (purple boxes). Due to the frameshift caused by the single nucleotide deletion in exon 5, the amino acid sequence of the Glass PA isoform gets shifted in *gl*^*del21.3*^ (orange box). B: Adult eye phenotype and expression of the retinal markers Rh1, NorpA, Trp, and Trpl of the *gl*^*del1.1*^ mutant. C: Adult eye phenotype and expression of the retinal markers Rh1, NorpA, Trp, and Trpl of the *gl*^*del22.4*^ mutant. D: Adult eye phenotype and expression of the retinal markers Rh1, NorpA, Trp, and Trpl of the *gl*^*del31.9*^ mutant. E: Adult eye phenotype and expression of the retinal markers Rh1, NorpA, Trp, and Trpl of the *gl*^*del21.3*^ mutant. C-E: All antibody stainings are shown in green, counterstaining of DNA with Hoechst (magenta). None of the tested photoreceptor makers is expressed in these glass alleles. Scale barrs: 40 µm F: ERGs of the different deletion alleles show no response to light, scale bars represent 5 mV (vertical) and 5 seconds (horizontal). G: Flies homozygous for the different small deletions in the *glass* locus are photoneutral. Two-tailed one sample *t* test followed by the Benjamini Hochberg procedure: For all data sets n = 10 experiments. *gl*^*del1.1*^: *p* = 0.90, *t*_*(9)*_ = 0.13; *gl*^*del21.3*^: *p* = 0.53, *t*_*(9)*_ = −1.11; *gl*^*del22.4*^: *p* = 0.49, *t*_*(9)*_ = −1.03; *gl*^*del31.9*^: *p* = 0.66, *t*_*(9)*_ = −0.57. The light preference index of all experimental groups is different from the light preference index of the control group shown in Figure 3. One-way ANOVA of preference indices: For all data sets n = 10 experiments, *p*<2*10^−16^, *F-Value*_*(8,81)*_= 44.04. CTRL vs *gl*^*del1.1*^: *p*<0.001, *t* = 8.74; CTRL vs *gl*^*del21.3*^: *p*<0.001, *t* = 9.81; CTRL vs *gl*^*del22.4*^: *p*<0.001, *t*=10.12; CTRL vs *gl*^*del31.9*^: *p*<0.001, *t* = 9.20. Data show mean and error bars show standard deviation. Red dots indicate means of individual experiments. ns = not significant.

