## Supplementary figures and images for "Multilevel regulation of the *glass* locus during *Drosophila* eye development"

### Suppl 1

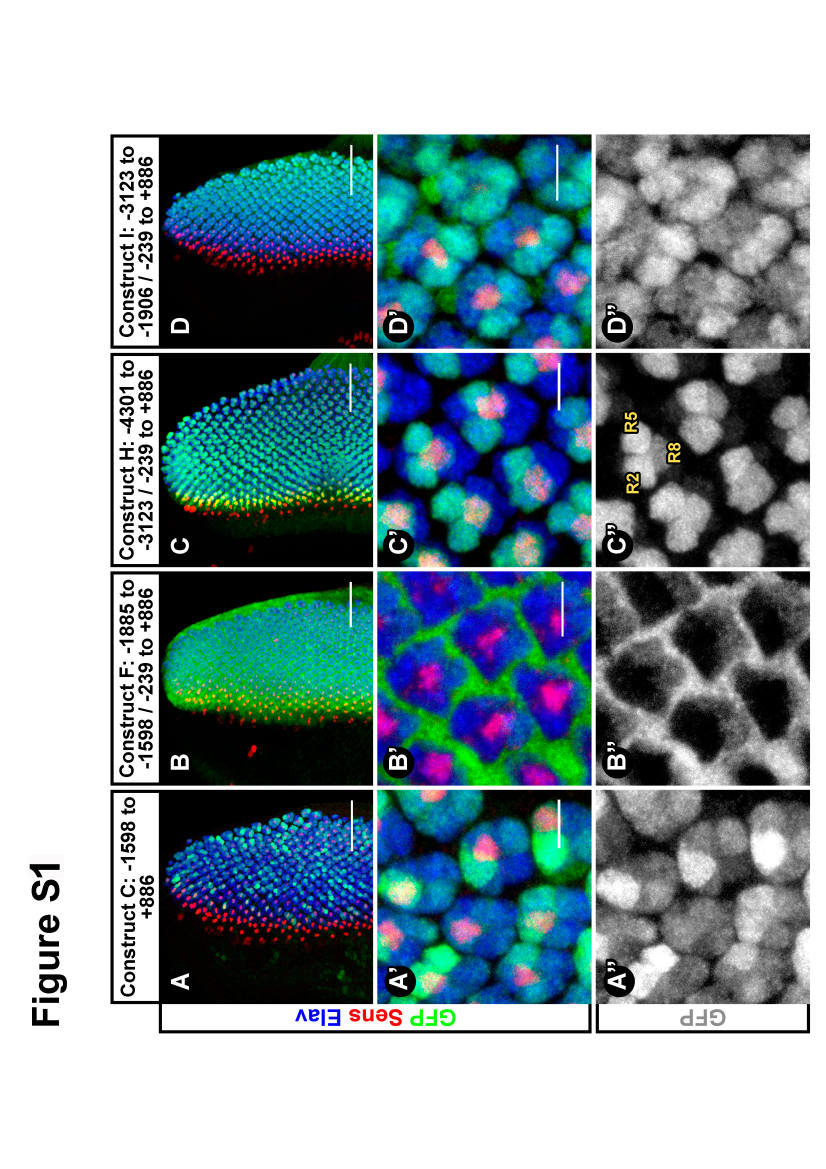

### Suppl 3

#### Figure S2

A

glass exon2 alignment

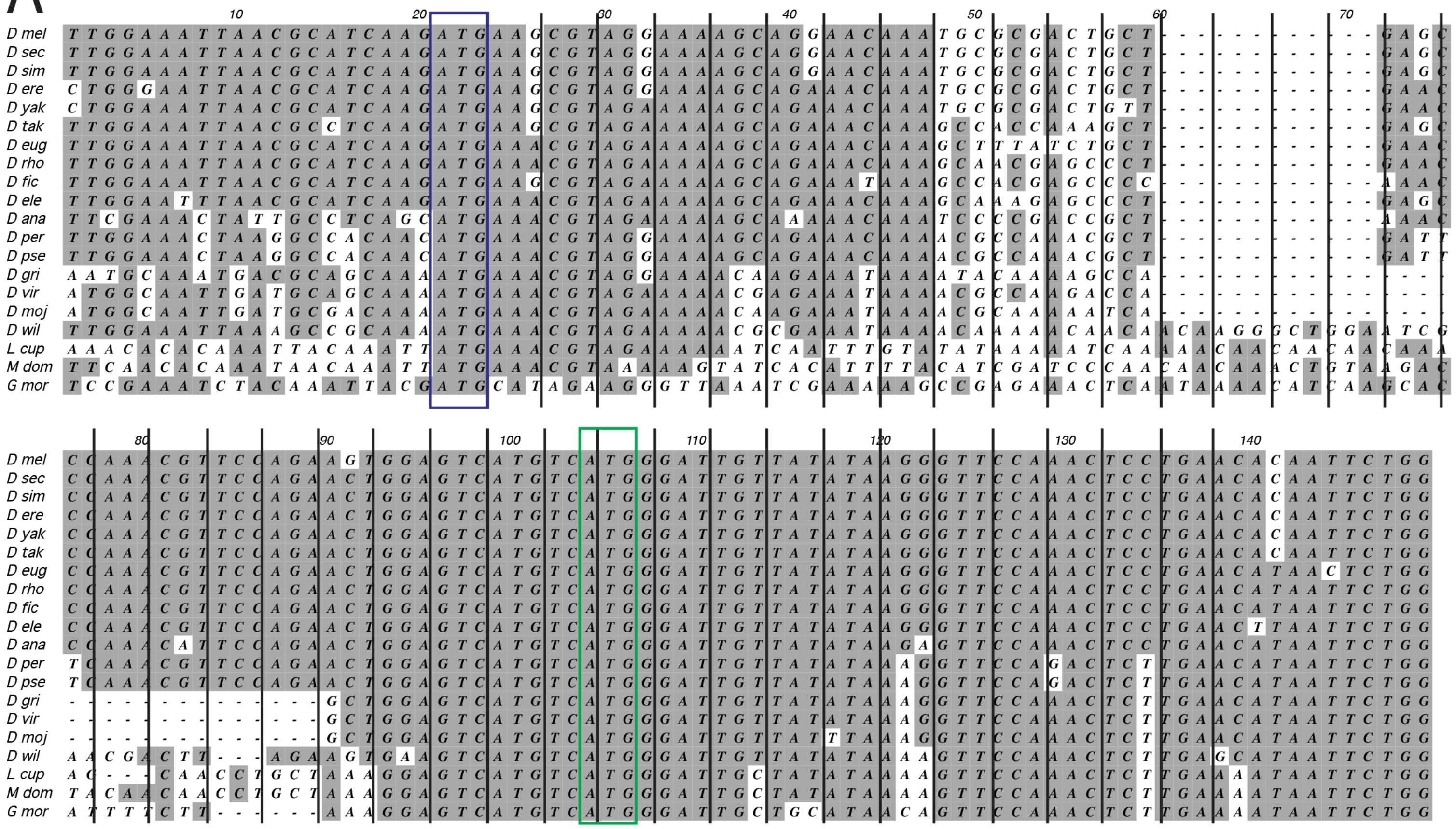

B

Dipteran vuORFs translated

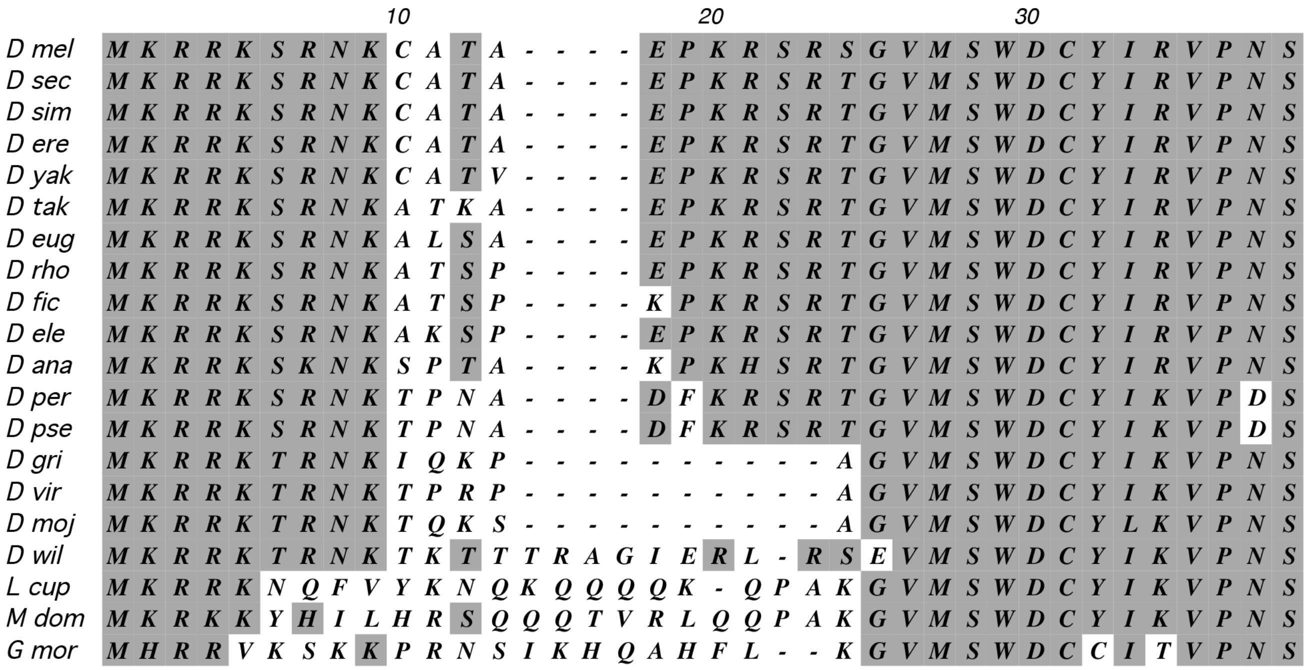

### Suppl 3

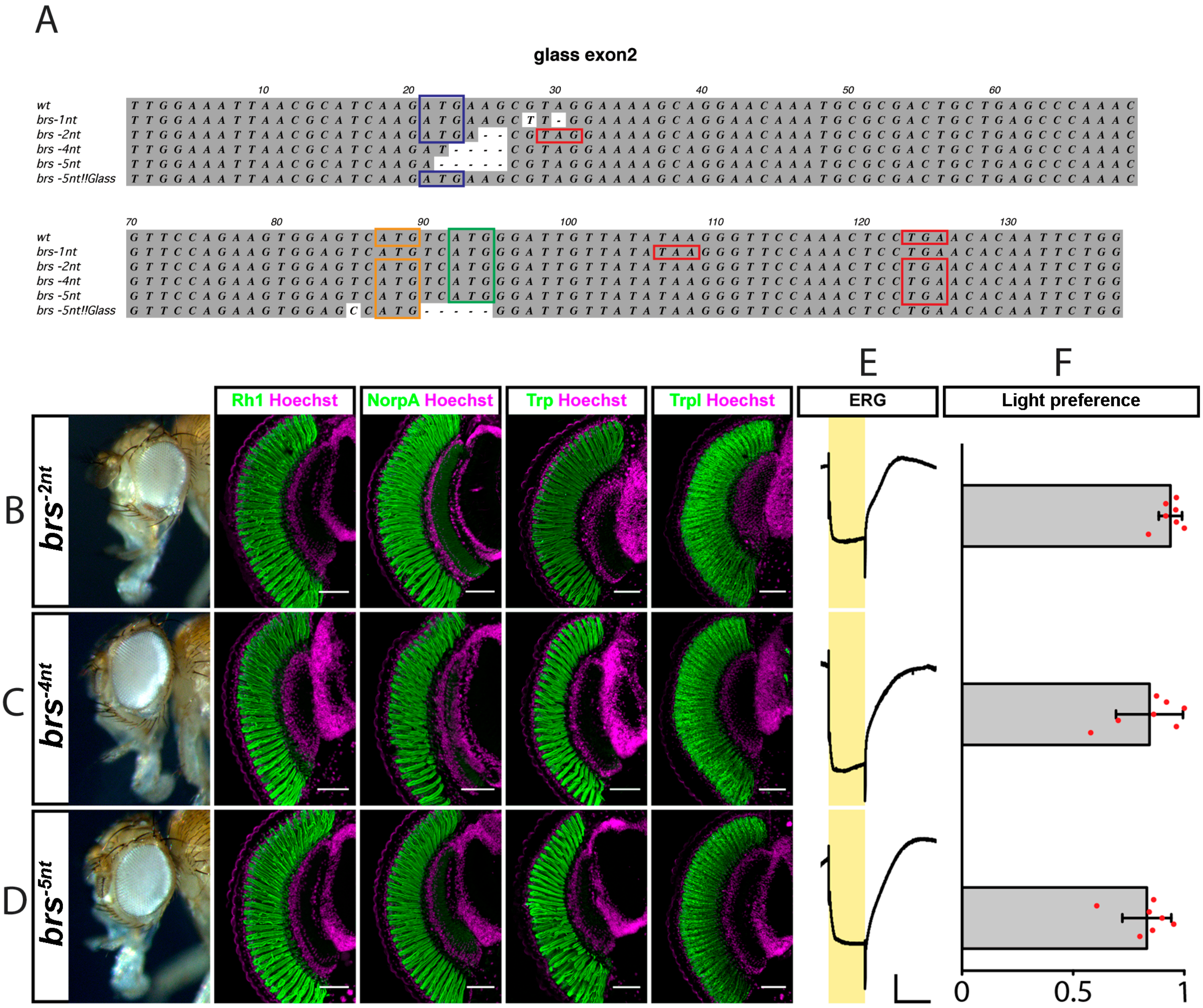

### Suppl 4

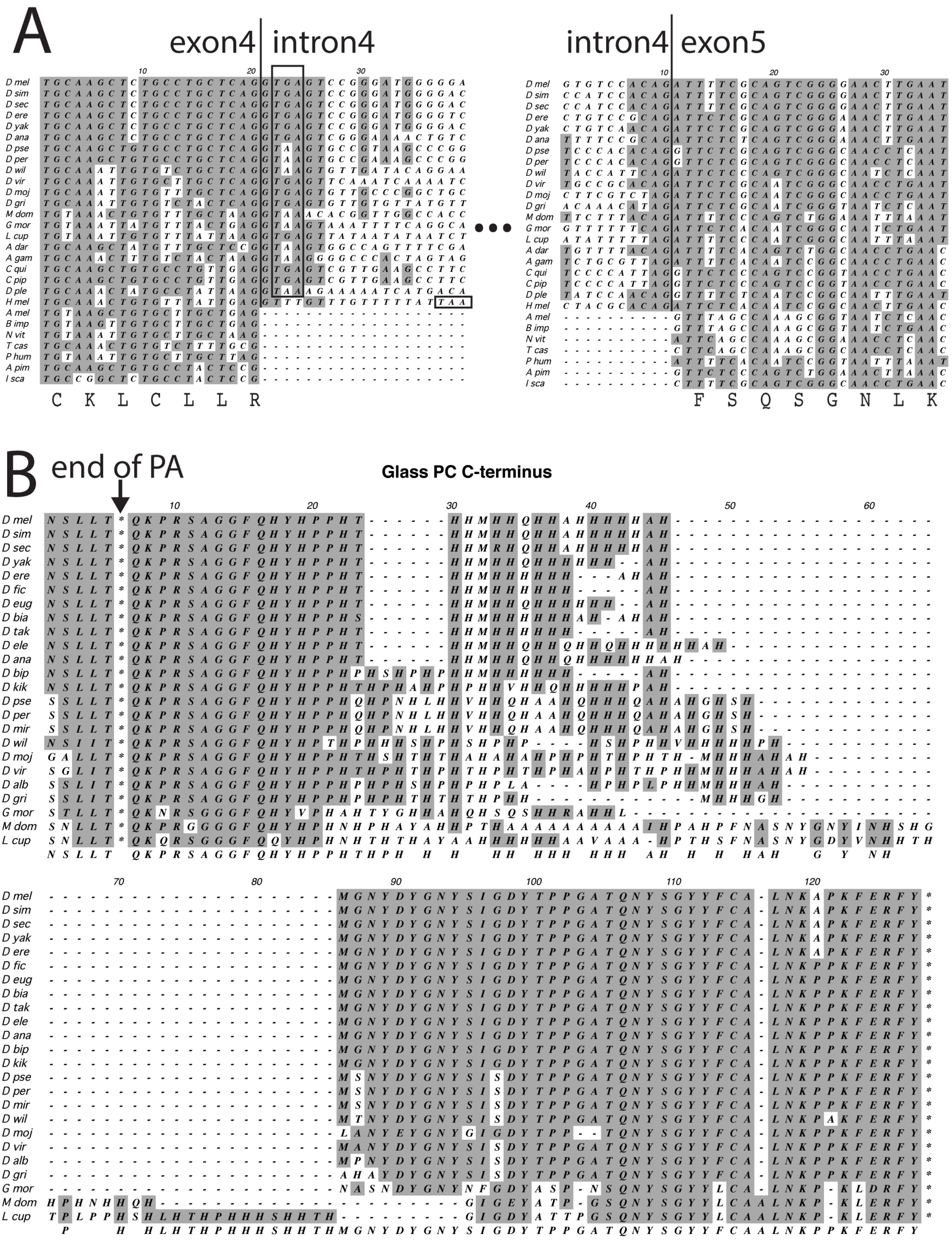

### Suppl 5

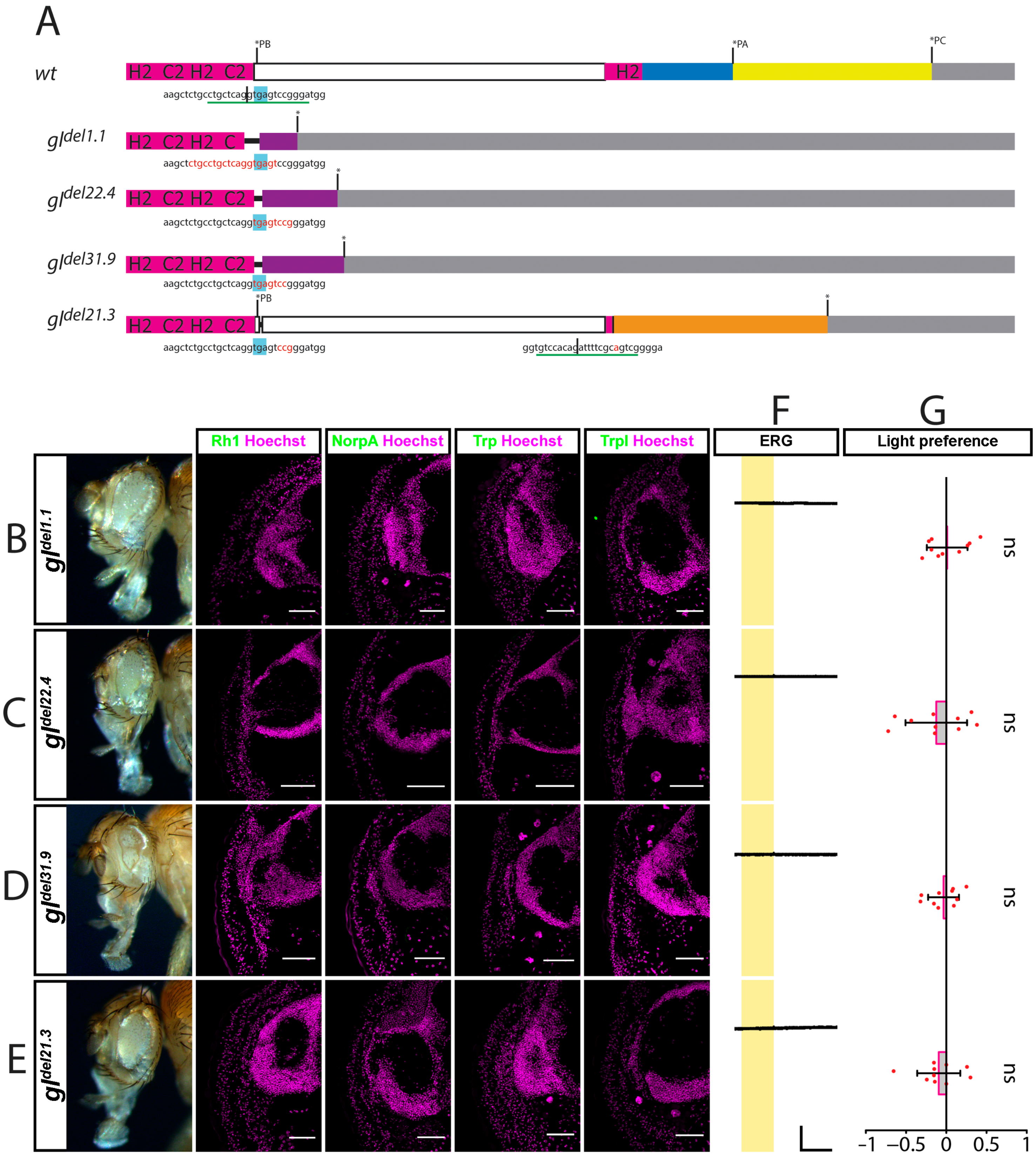
